# Hidden in plain sight: Discovery of sand flies in Singapore and description of four species new to science

**DOI:** 10.1101/2025.02.06.636818

**Authors:** Huicong Ding, Majhalia Torno, Khamsing Vongphayloth, Germaine Ng, Denise Tan, Wendy Sng, Kelvin Ho, Fano José Randrianambinintsoa, Jérôme Depaquit, Cheong Huat Tan

## Abstract

Phlebotomine sand flies (Diptera: Psychodidae) are small, blood-sucking insects that are of significant public and veterinary health importance for their role in the transmission of *Leishmania* parasites, bacteria and arboviruses. Although sand flies have been documented in most Southeast Asian countries, there are no published records confirming their presence in Singapore. Here, we provide the first documented evidence on the presence of sand flies in Singapore. Using an integrated taxonomic approach that combines morphological analysis with DNA barcoding of the mitochondrial *cytochrome b* (*cytb*) and *cytochrome c oxidase subunit I* (*COI*) genes, we identified eight sand fly species, including four newly described species – *Phlebotomus seowpohi* n. sp., *Sergentomyia leechingae* n. sp., *Sergentomyia gubleri* n. sp., and *Sergentomyia retrocalcarae* n. sp.. Phylogenetic analyses suggest that the new *Phlebotomus* species, belonging to subgenus *Euphlebotomus*, is closely related to *Phlebotomus argentipes*, an important vector of *Leishmania donovani* from the South Asian region. The discovery of phlebotomine sand flies in Singapore underscores the importance of biosurveillance of biting arthropods. Given Singapore’s status as a major travel hub, there is a potential risk of leishmaniasis being introduced either by residents returning, or visitors arriving from endemic regions. This risk is compounded by the recent detection of local canine leishmaniasis. Thus, continuous monitoring is essential to assess and manage the risk of disease transmission, support the development of an early warning system, and enable timely and targeted public health interventions. The findings from this study contributes to the global knowledge on sand fly and enhance our understanding of local sand fly diversity and distribution.

**Author Summary:** Phlebotomine sand flies are small, blood-sucking insects recognized as significant vectors for various diseases including leishmaniasis. This study provides the first evidence of their presence in Singapore. Using an integrated taxonomic approach combining morphological analysis with DNA barcoding, we identified eight sand fly species that include four new species: *Phlebotomus seowpohi* n. sp., *Sergentomyia leechingae* n. sp., *Sergentomyia gubleri* n. sp., and *Sergentomyia retrocalcarae* n. sp. The presence of sand flies in Singapore highlights an important biosurveillance need, as the region’s status as a major travel hub increases the risk of leishmaniasis being introduced. Importantly, recent findings of canine leishmaniasis in local dogs amplify the threat of disease transmission. Continuous monitoring and understanding of sand fly diversity and ecology are essential for developing effective public health strategies and early warning systems to manage this risk.

## 1. Introduction

Phlebotomine sand flies are small (2 – 3 mm in length), hematophagous insects belonging to the order Diptera under the family Psychodidae. They are of significant public and veterinary health importance, with many members of this group incriminated as vectors of viral, bacterial, and parasitic diseases [1,2]. They are widely known for their role in the transmission of leishmaniasis, a neglected tropical disease (NTD) caused by an obligate, intracellular protozoan parasite belonging to the genus *Leishmania* [3,4].

Leishmaniasis manifests in many forms, with visceral leishmaniasis being the most significant and severe form. If left untreated, it is fatal in over 95% of cases [5]. As with most NTDs, people living in impoverished communities are disproportionately affected, with reported cases primarily concentrated in the Americas, East Africa, Middle East, and Central Asia [6]. Despite its high morbidity, management of leishmaniasis is hindered by difficulty in managing vector population, absence of vaccines, inaccurate disease diagnosis, and ineffective treatment options [7].

More than a thousand phlebotomine sand fly species have been described globally [8], with approximately 80 species identified to be vectors of leishmaniasis [9]. However, there is limited information about sand flies and their diversity in Southeast Asia (SEA), this is largely due to the region’s long-standing history of being “leishmaniasis-free”. The detection of the first autochthonous visceral leishmaniasis case in Thailand in 1996 led to increased interest in the study of the region’s sand flies [10]. In recent years, additional reports of indigenous leishmaniasis cases due to *Leishmania martiniquensis* and *L. orientalis* in different provinces of Thailand underscore the need for a comprehensive approach in understanding the epidemiology of leishmaniasis, as well as the biology and diversity of sand flies in the region [11,12].

Most of the research on phlebotomine sand flies in SEA is concentrated in Thailand [13–25], with occasional surveys conducted in Cambodia [26], Indonesia [27], Laos [13,28,29], Malaysia [30–32], Philippines [33–35], and Vietnam [23,36]. Currently, the region records about 60 sand fly species, belonging to the following genera: *Phlebotomus*, *Sergentomyia*, *Grassomyia*, *Idiophlebotomus,* and *Chinius* [13,22,37]. This is likely to be underestimated due to limited biosurveillance efforts being carried out in the region.

Singapore (1.3521° N, 103.8198° E) is an island state located at the southern tip of the Malay Peninsula. Despite the tropical climate being an ideal habitat for sand flies [2,38–43], to the best of our knowledge, there are no official records of phlebotomine sand flies in Singapore. Here, we report for the first time the presence of sand flies in Singapore. Additionally, we formally describe four newly identified species using an integrated taxonomy approach involving morphology analyses and DNA barcoding: *Phlebotomus (Euphlebotomus) seowpohi* n. sp., *Sergentomyia gubleri* n. sp., *Sergentomyia leechingae* n. sp., and *Sergentomyia retrocalcarae* n. sp. The species descriptions include illustrations of their head, genitalia, and wing, along with DNA barcodes of the mitochondrial *cytochrome b* (*cytb*) and *cytochrome c oxidase subunit I* (*COI*) gene fragments. The findings from this study will contribute to a comprehensive global sand fly diversity checklist, which is essential for advancing sand fly research. Locally, a comprehensive understanding of sand fly diversity and distribution can provide valuable insight into environmental factors that determines where they are most likely to thrive. This knowledge can be useful for developing risk maps that can help to identify potential sand fly hot spot and help guide the development of targeted vector control strategy to manage sand fly population.

## 2. Methods

### Study areas

Singapore has a total land area of 742.22 km^2^, with 49% vegetation cover [44]. The remaining 51% is of highly urbanized landscape consisting of buildings and other artificial impervious surfaces. Situated one degree north of the equator, Singapore has a tropical rainforest climate with abundant rainfall and uniformly high temperatures year-round with no distinct seasons [45]. The average daily temperature ranges between 23°C - 33°C with an average annual rainfall of 2165.9 mm.

The entomological surveillance period was from September 2020 to September 2021 at 29 sampling sites across Singapore (S1 Fig). These sites are part of Singapore’s One Health biosurveillance effort for vector-borne diseases, and include secondary growth forests, public parks, fringes of nature reserves, peri-urban vegetated areas, open fields, coastal areas, and urban areas.

### Sand fly collection

Night Catcher (NC) traps (Orinno Technology Pte Ltd, Singapore) were used to collect the sand flies (Fig 1). These are modified CDC light traps with the additional feature of a rotating module that separates collections at hourly intervals from 1900 h to 0700 h, to coincide with most sand flies peak biting time [46,47]. At each sampling site, two to eight NC traps were placed at least 50m apart. Dry ice, to generate a slow release of carbon dioxide, and incandescent lights were used as trap attractants. The GPS coordinates of each trap location were recorded. Collection bottles containing captured specimens were transported to the laboratory and placed in a −20°C freezer overnight. After freezing, specimens were sorted, and sand flies were stored in 80% ethanol prior to species identification.

**Figure 1.**
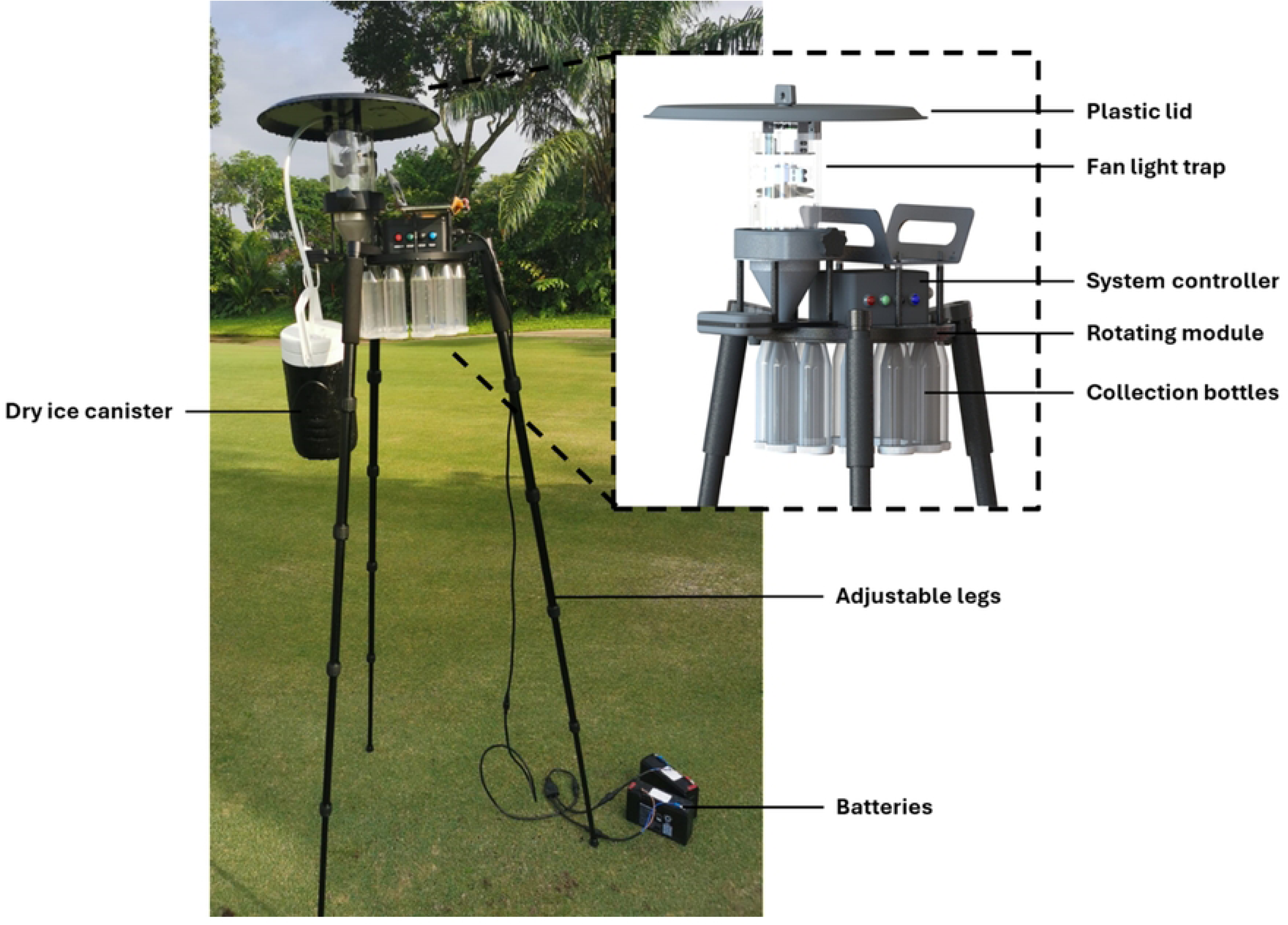
Night Catcher trap deployed for sand fly collection.

### Sample processing

Specimens were initially sorted to morpho-groups under *Phlebotomus* and *Sergentomyia*, using available keys for international [48,49], as well as Southeast Asian phlebotomine sand flies [23,35,50–53]. For each sampling location within a morpho-group, at least one male and one female specimen was subsampled for molecular analyses. The subsampled specimens were subjected to a non-destructive DNA extraction protocol and vouchered. After which, morphological analyses were performed to clarify taxonomic affinities.

### Non-destructive DNA extraction

Non-destructive DNA extraction was performed individually using the DNeasy Blood and Tissue Kit (Qiagen, Hilden, Germany), retaining morphological integrity of each specimen while extracting sufficient DNA material for downstream sequencing. In this process, sand fly specimens stored in 80% ethanol were rinsed three times with ultrapure water to remove excess ethanol. To minimize damage to the specimens during rinsing, water was removed and replaced using a pipette, instead of moving the specimens into new tubes. After rinsing, 180 µL Buffer ATL and 20 µL Proteinase K mixture were added to each specimen. This was followed by whole-specimen incubation at 56 °C for 1 hr. No dissection or tissue homogenization was involved. After incubation, DNA extraction followed the manufacturer’s standard DNA extraction protocol with slight modifications – genomic DNA was eluted twice, first in 50 µL Buffer AE, then in 100 µL Buffer AE. The remaining sand fly carcass was rinsed once with ultrapure water to remove all buffer residues before preservation in 80% ethanol for subsequent morphological analyses.

### Morphological analyses

Soft tissue was further lysed in a bath of 10% KOH, then bleached in Marc-André solution. Bleached specimens were subjected to a series of alcohol baths for dehydration, then mounted onto microscope slides with Euparal^®^ and covered with cover slips for species identification. Visual analysis of the specimens was performed using an Olympus BX61 compound microscope (Olympus, Japan). Measurements and counts were made by using the Stream Motion software (Olympus, Japan) with a video camera connected to the microscope. For measurements of curved parts, the “polygonal line” tracing tool was used to acquire the most accurate measurement. Drawings were made using an Olympus camera lucida.

Specimens were analyzed using the previously mentioned available keys. The terminologies and abbreviations adopted for the morphological characters of phlebotomine sand flies follow that of Galati et al. (2017) [54].

### Molecular analyses

Polymerase chain reaction (PCR) amplification was performed in a 50 µL volume using 5 µL of extracted DNA and 50 pmol each of forward and reverse primers. The PCR mix contained (final concentrations): 10 mM Tris HCl (pH 8.3), 1.5 mM MgCl2, 50 mM KCl, 0.01% Triton X-100, 200 µM dNTP for each base, and 1.25 units of 5 prime Taq polymerase (Eppendorf, Germany). PCR amplification was undertaken using the primers C3B-PDR (5’-CAYATTCAACCWGAATGATA-3’) and N1N-PDR (5’- GGTAYWTTGCCTCGAWTTCGWTATGA-3’) for *cytb* [55], and LCO1490 (5’- GGTCAACAAATCATAAAGATATTGG-3’) and HCO2198 (5’- TAAACTTCAGGGTGACCAAAAAATCA-3’) for *COI* gene fragments [56]. These amplifications were performed according to conditions published previously [55,56]. Amplicons were purified using the E-Gel Clonewell II gel (Invitrogen, USA) and sent for sequencing in both directions using the same primers.

Molecular analyses were performed on consensus *cytb* and *COI* sequences, respectively. Multiple alignment of consensus sequences was carried out using the ClustalW algorithm [57], in software MEGA 11 [58]. This was followed by the construction of a maximum likelihood (ML) tree with 100 bootstrap replicates. Construction was based on the substitution model selected by Modeltest [59] with an Akaike information criterion (AIC) of HKY85 [60]. Available *cytb* and *COI* sequences belonging to other known sand fly species were retrieved from GenBank [61] and included in the construction of the ML tree. Inter- and intra-specific pairwise distances were also calculated using MEGA 11 [58].

## 3. Results

### 3.1 Inventory

A total of 223 phlebotomine sand flies were collected over a one-year period (Table 1). Sand flies were collected at 15 of the 29 sampling sites (Fig 2). Of these, 127 (57%) were identified as females and the remaining 43% were males. The collected sand flies represented eight species across two genera, *Phlebotomus* (n=192, 86.1%) and *Sergentomyia* (n=31, 13.9%). These species are *Ph. stantoni*, *Se. barraudi* group, *Se. iyengari* group, *Se. whartoni*, and the newly described *Ph. seowpohi* n. sp., *Se. gubleri* n. sp., *Se. leechingae* n. sp., and *Se. retrocalcarae* n. sp.. The new species were described through an integrative taxonomy approach using morphological and molecular methods.

**Fig 2.**
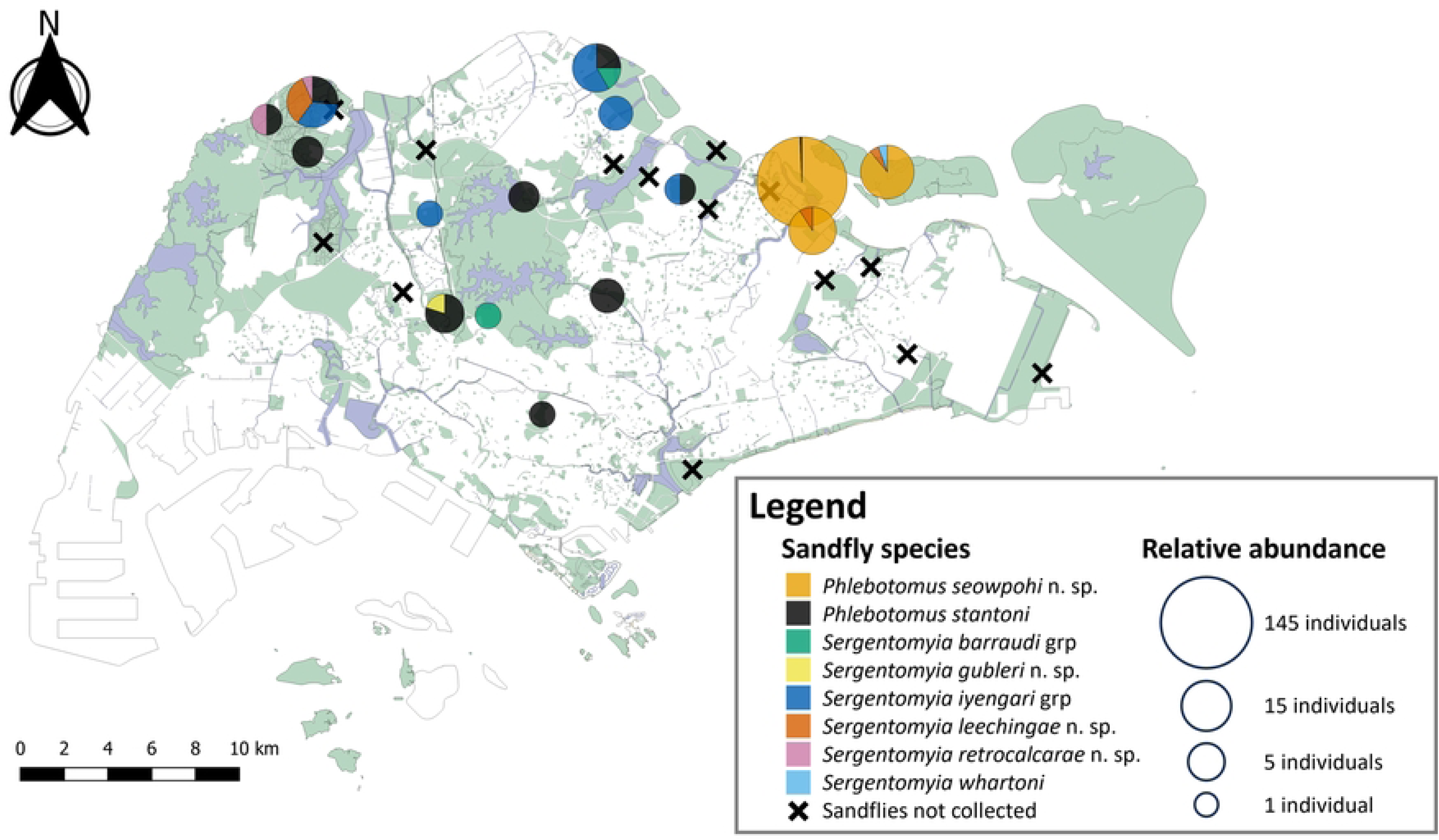
Sand fly species composition across fifteen positive sampling sites in Singapore. Pie charts are scaled to the square root of the total number of individuals caught at each sampling site. Fourteen sampling sites where no sand flies were collected during this study are indicated with an ×. A total of twenty-nine sampling sites were sampled in this study.

**Table 1.**
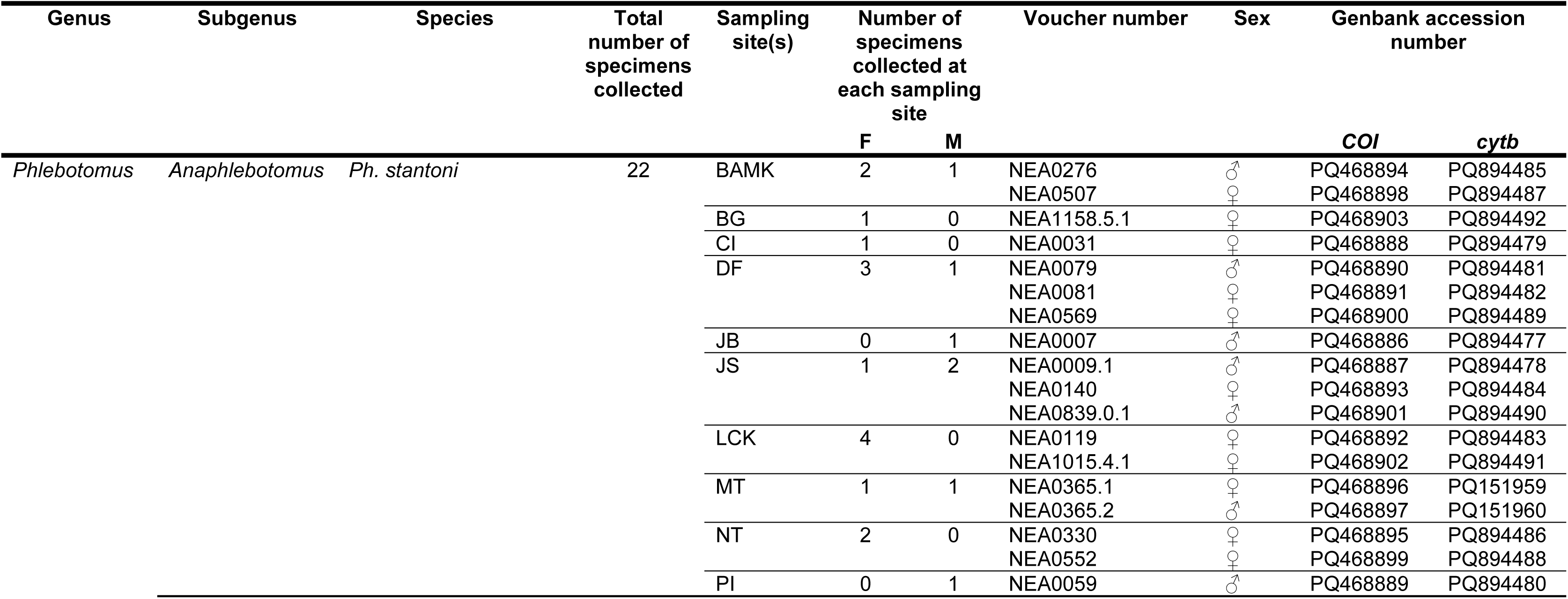
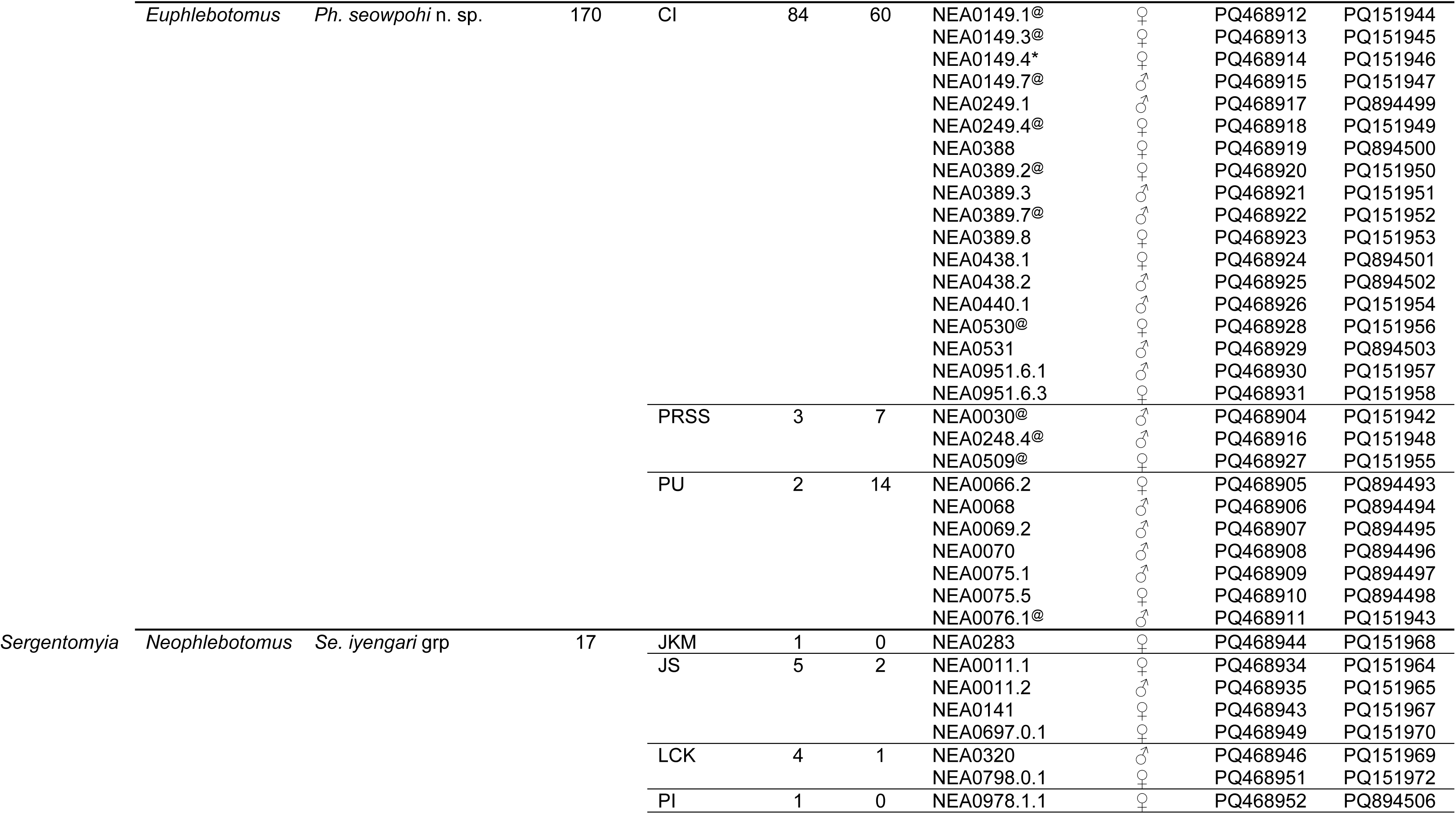
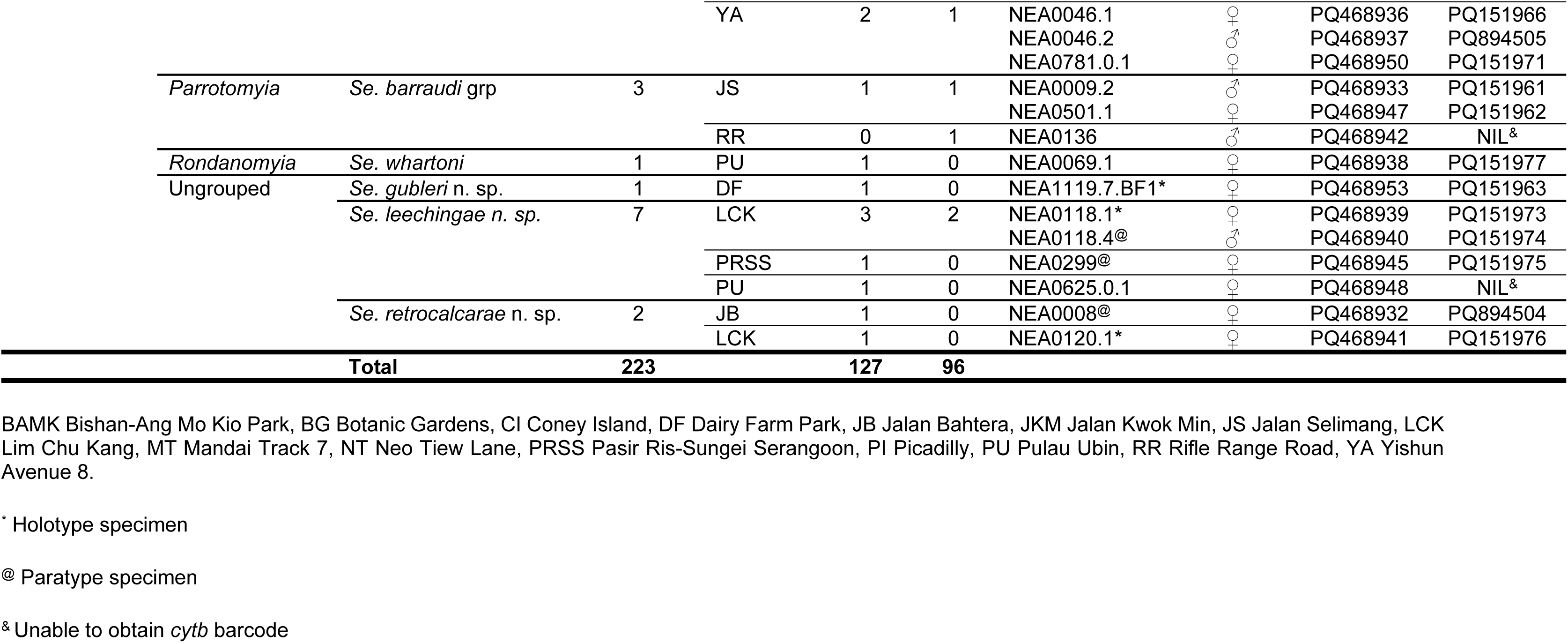
Sand fly species collected in Singapore from September 2020 to September 2021 Specimen voucher numbers and Genbank accession numbers for c*ytb* and *COI* sequences generated in this study are listed.

The majority (n=170, 76.2%) of the specimens caught were identified as *Ph. Seowpohi* n. sp., but its distribution appears to be limited to the coastal regions of northeastern Singapore (Fig 2).

### 3.2 Molecular analyses of phlebotomine specimens

Altogether, 68 sand fly specimens were successfully barcoded (Table 1). Of which, 66 specimens had both their *cytb* and *COI* gene fragments amplified and sequenced. Consensus *cytb* sequences were 411 bp and 410 bp for *Phlebotomus* and *Sergentomyia* specimens, respectively. These consensus sequences were used to generate maximum likelihood phylogenetic trees for *Phlebotomus* (Fig 3) and *Sergentomyia* (Fig 4) species. Sand fly species collected in Singapore formed clades with high bootstrap support (>92%) with the exception of the single sequences representing *Se. gubleri* n. sp. and *Se. whartoni*. Intraspecific *cytb* p-distance was less than 3.3% for species found in Singapore, while interspecific p-distance ranged from 12.5% to 21.3% when compared with reference barcodes. The calculations of p-distance over sequence pairs between and within *Phlebotomus* and *Sergentomyia* species are provided in Table 2 and Table 3, respectively.

**Fig 3.**
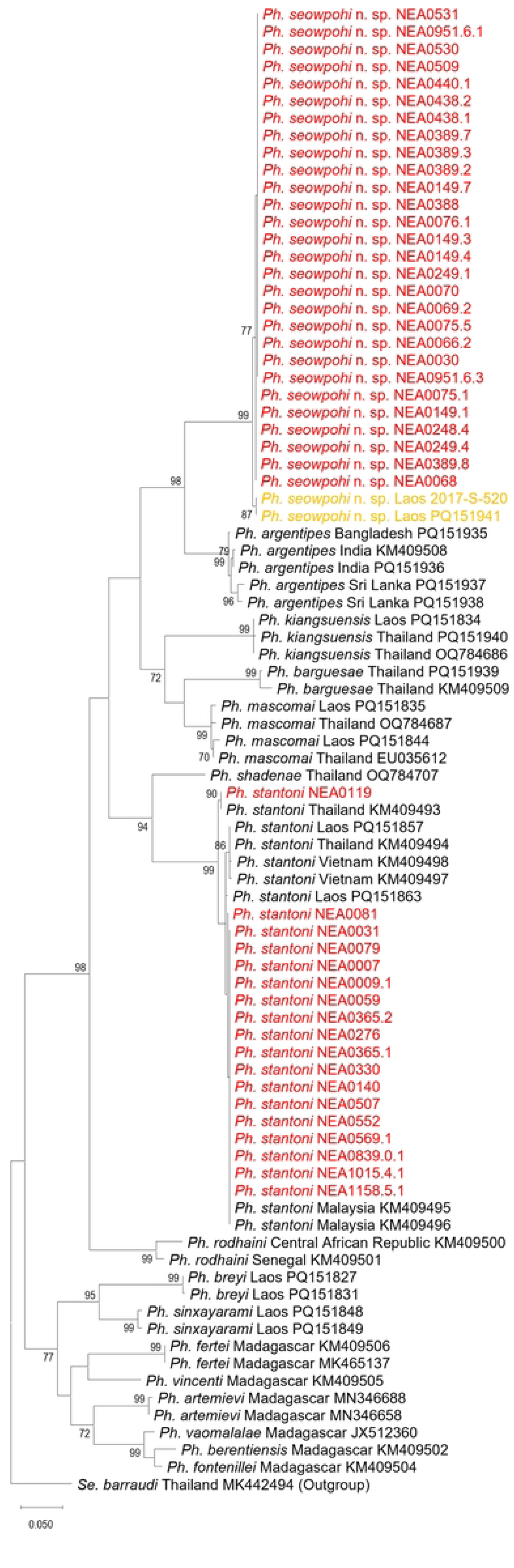
Maximum-likelihood *Phlebotomus* phylogenetic tree inferred from aligned consensus *cytb* sequences using the Hasegawa-Kishino-Yano 85 model. Sequences highlighted in red are generated from sand fly specimens collected in Singapore. Sequences in yellow are from Khammouane and Luangphabang, Laos, as part of IP Laos collection program. They will be further detailed in the discussion section. The trees have been rooted on a reference *Sergentomyia barraudi* sequence which is selected as an outgroup. Nodes with bootstrap value less than 70% are not shown.

**Fig 4.**
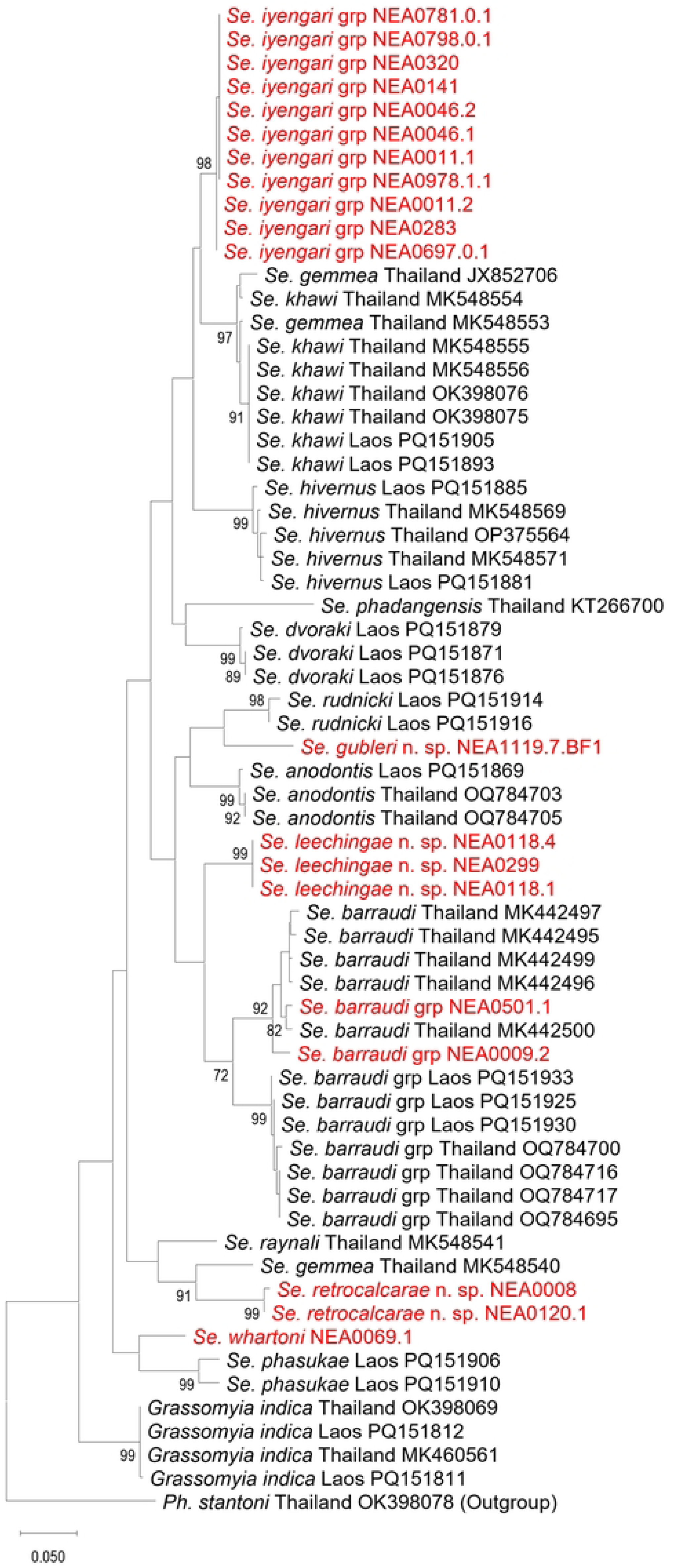
Maximum-likelihood *Sergentomyia* phylogenetic tree inferred from aligned consensus *cytb* sequences using the Hasegawa-Kishino-Yano 85 model. Sequences highlighted in red are generated from sand fly specimens collected in Singapore. The trees have been rooted on a reference *Phlebotomus stantoni* sequence which is selected as an outgroup. Nodes with bootstrap value less than 70% are not shown.

**Table 2.** Mean interspecific genetic distances for *cytb* sequence pairs between five *Phlebotomus (Euphlebotomus)* species and *Ph. (Anaphlebotomus) stantoni*. Numbers (1-8) in the top row correspond to the species listed. Calculations were based on the p-distance model. Diagonal bold values indicate intraspecific mean distances. Names with an asterisk refer to specimens collected in Singapore (SG).

|  | Species | 1 | 2 | 3 | 4 | 5 | 6 | 7 | 8 |
| --- | --- | --- | --- | --- | --- | --- | --- | --- | --- |
| 1 | *Ph. seowpohi* n. sp.* | **0.001** |  |  |  |  |  |  |  |
| 2 | *Ph. stantoni* * | 0.211 | **0.002** |  |  |  |  |  |  |
| 3 | *Ph. argentipes* | 0.125 | 0.197 | **0.014** |  |  |  |  |  |
| 4 | *Ph. barguesae* | 0.200 | 0.212 | 0.183 | **0.016** |  |  |  |  |
| 5 | *Ph. kiangsuensis* | 0.182 | 0.207 | 0.173 | 0.169 | **0.002** |  |  |  |
| 6 | *Ph. mascomai* | 0.159 | 0.171 | 0.143 | 0.117 | 0.141 | **0.010** |  |  |
| 7 | *Ph. seowpohi* n. sp. (Laos) | 0.010 | 0.213 | 0.125 | 0.202 | 0.180 | 0.161 | **0** |  |
| 8 | *Ph. stantoni* (non-SG) | 0.210 | 0.009 | 0.198 | 0.213 | 0.207 | 0.171 | 0.212 | **0.010** |

**Table 3.** Mean interspecific genetic distances for *cytb* sequence pairs between *Sergentomyia* species and *Grassomyia* species. Numbers (1-17) in the top row correspond to the species listed. Calculations were based on the p-distance model. Diagonal bold values indicate intraspecific mean distances. Names with an asterisk refer to specimens collected in Singapore (SG).

|  | Species | 1 | 2 | 3 | 4 | 5 | 6 | 7 | 8 | 9 | 10 | 11 | 12 | 13 | 14 | 15 | 16 | 17 |
| --- | --- | --- | --- | --- | --- | --- | --- | --- | --- | --- | --- | --- | --- | --- | --- | --- | --- | --- |
| 1 | *Se. barraudi* grp* | **0.033** |  |  |  |  |  |  |  |  |  |  |  |  |  |  |  |  |
| 2 | *Se. gubleri* n. sp.* | 0.140 | **NA** |  |  |  |  |  |  |  |  |  |  |  |  |  |  |  |
| 3 | *Se. iyengari* grp* | 0.109 | 0.123 | **0.001** |  |  |  |  |  |  |  |  |  |  |  |  |  |  |
| 4 | *Se. leechingae* n. sp.* | 0.108 | 0.120 | 0.105 | **0** |  |  |  |  |  |  |  |  |  |  |  |  |  |
| 5 | *Se. retrocalcarae* n. sp.* | 0.148 | 0.136 | 0.136 | 0.133 | **0.005** |  |  |  |  |  |  |  |  |  |  |  |  |
| 6 | *Se. whartoni* * | 0.131 | 0.120 | 0.096 | 0.132 | 0.124 | **NA** |  |  |  |  |  |  |  |  |  |  |  |
| 7 | *Grassomyia indica* | 0.143 | 0.147 | 0.129 | 0.142 | 0.139 | 0.113 | **0.001** |  |  |  |  |  |  |  |  |  |  |
| 8 | *Se. anodontis* | 0.119 | 0.114 | 0.094 | 0.094 | 0.149 | 0.125 | 0.134 | **0.005** |  |  |  |  |  |  |  |  |  |
| 9 | *Se. barraudi* grp | 0.057 | 0.125 | 0.117 | 0.091 | 0.157 | 0.134 | 0.143 | 0.104 | **0.044** |  |  |  |  |  |  |  |  |
| 10 | *Se. dvoraki* | 0.125 | 0.127 | 0.084 | 0.118 | 0.136 | 0.120 | 0.146 | 0.131 | 0.135 | **0.005** |  |  |  |  |  |  |  |
| 11 | *Se. gemmea* | 0.132 | 0.114 | 0.079 | 0.115 | 0.131 | 0.117 | 0.133 | 0.119 | 0.134 | 0.115 | **0.093** |  |  |  |  |  |  |
| 12 | *Se. hivernus* | 0.116 | 0.122 | 0.073 | 0.121 | 0.134 | 0.099 | 0.129 | 0.123 | 0.127 | 0.109 | 0.106 | **0.010** |  |  |  |  |  |
| 13 | *Se. khawi* | 0.127 | 0.099 | 0.050 | 0.103 | 0.142 | 0.106 | 0.130 | 0.102 | 0.122 | 0.106 | 0.052 | 0.088 | **0.004** |  |  |  |  |
| 14 | *Se. phadangensis* | 0.121 | 0.138 | 0.127 | 0.164 | 0.154 | 0.128 | 0.147 | 0.142 | 0.131 | 0.140 | 0.142 | 0.129 | 0.125 | **NA** |  |  |  |
| 15 | *Se. phasukae* | 0.131 | 0.132 | 0.119 | 0.132 | 0.146 | 0.097 | 0.132 | 0.139 | 0.130 | 0.140 | 0.125 | 0.126 | 0.111 | 0.142 | **0.035** |  |  |
| 16 | *Se. rudnicki* | 0.114 | 0.095 | 0.114 | 0.108 | 0.142 | 0.103 | 0.142 | 0.098 | 0.106 | 0.131 | 0.127 | 0.132 | 0.114 | 0.140 | 0.116 | **0.010** |  |
| 17 | *Se. raynali* | 0.117 | 0.140 | 0.121 | 0.119 | 0.116 | 0.119 | 0.122 | 0.126 | 0.128 | 0.134 | 0.117 | 0.127 | 0.118 | 0.152 | 0.114 | 0.104 | **NA** |

NA denotes cases in which it was not possible to estimate genetic distances due to single DNA barcode.

Similarly, *COI* sequences generated are aligned and trimmed to a final length of 493 bp and 536 bp for *Phlebotomus* and *Sergentomyia* specimens respectively. Results of molecular analyses on consensus *COI* sequences are comparable to that of the *cytb* sequences. These are provided in S2 Fig and S1 Table.

### 3.3 Description of new species

#### 3.3.1 Description of *Phlebotomus seowpohi* Depaquit, Vongphayloth, and Tan n. sp

Genus *Phlebotomus* Rondani & Berté, in Rondani 1840.

Subgenus *Euphlebotomus*, Theodor 1948.

The coloration of *Ph. seowpohi* n. sp. is brown, with a dark brown scutum, in both sexes.

##### Etymology

The epithet *seowpohi* refers to Mr. Seow-Poh Khoo, former Director General and Deputy Chief Executive Officer of the National Environment Agency (NEA) Singapore, in recognition of his significant contributions to improve Singapore’s public health including prevention and control of vector-borne diseases.

Type locality: Coney Island (1°24’31.5”N 103°55’20.9”E), Singapore

##### Type specimens

Female holotype (voucher NEA0149.4), two female paratypes (voucher NEA0249.4 and NEA0530), and two male paratypes (voucher NEA0149.7, NEA0030) will be deposited at the Laboratory of Entomology, The Natural History Museum, London, UK.

Three female paratypes (voucher NEA0149.1, NEA0149.3, and NEA0509) and two male paratypes (voucher NEA0076.1 and NEA0248.4) deposited at the Environmental Health Institute, National Environment Agency, Singapore.

One female paratype (voucher NEA0389.2) and one male paratype (voucher NEA0389.7) will be deposited at the Lee Kong Chian Natural History Museum, Singapore.

To meet the criteria of availability, the authors Depaquit, Vongphayloth, and Tan are responsible for the name *Phlebotomus seowpohi* n. sp. and should be cited as the sole authority of this taxa, according to the Article 50 (1) of the lnternational Code of Zoological Nomenclature.

###### 3.3.1.1 Description of the female of *Ph. seowpohi* n. sp. (Fig 5)

Description and measurements indicated are those of the female holotype labelled NEA0149.4. Measurements carried out on several specimens are indicated in S3 Table.

**Figure 5.**
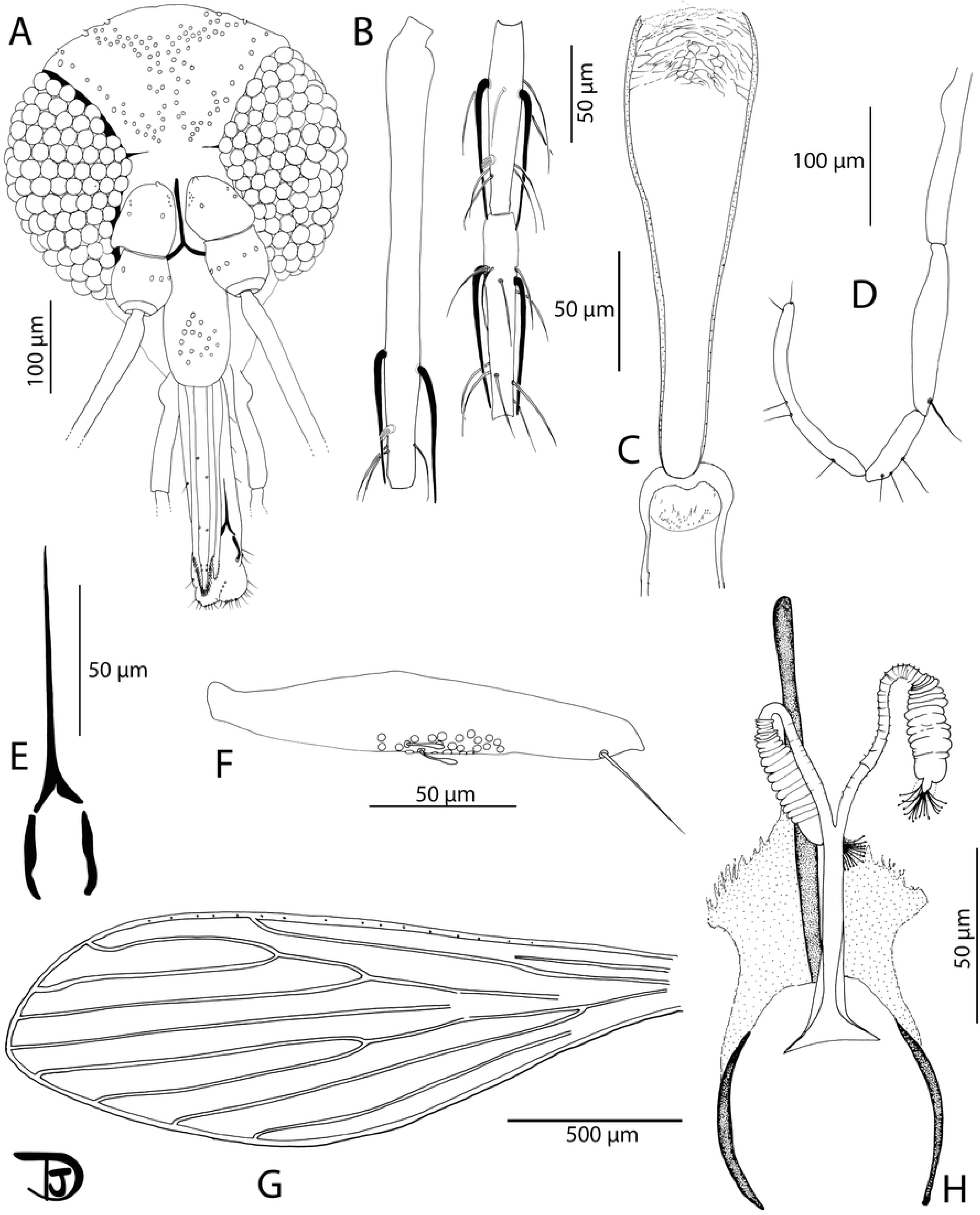
*Phlebotomus seowpohi* n. sp. female. A: head; B: flagellomeres 1, 2, and 3; C: pharynx and cibarium; D: palpi; E: labial furca; F: 3rd palpal article; G: wing; H: genital furca and spermathecae.

###### 3.3.1.1.1 Head

Occiput with two narrow lines of well individualised setae. Clypeus: 155 µm long, 75 µm wide, exhibiting 11 setae. Eyes: 270 µm long, 145 µm wide, with about 120 facets. Incomplete interocular sutures. Flagellomeres: f1 (=AIII) = 247 µm, f2 (=AIV) = 106 µm, f3 (=AV) = 104 µm. Flagellomere 1 longer than f2+f3. Presence of two long ascoids from f1 to f13, not reaching the next articulation, except those of f1 (observed on paratype NEA0149.3). Ascoid of f2: 74.8 µm long. Length of this ascoid/length of f2 = 0.71. One distal papilla on f1 and f2. Absence of papillae on f3 to f8. Two papillae on f9, four on f10, three on f11, five on f12 and f13, and four on f14. Several simple setae on all flagellomeres. Palps: p1 = 35 µm, p2 = 104 µm, p3 = 147 µm, p4 = 69 µm, p5 = 186 µm. Palpal formula: 1, 4, 2, 3, 5. Presence of one distal spiniform seta on p3, four on p4, and six on p5. Labrum: 256 µm long. f1/labrum = 0.96. Lateral teeth of the hypopharynx not observable. Teeth of the maxillary lacinia not observable. Labial furca closed. Cibarium armed with 5-6 anterior teeth pointed backwards with 30 lateral teeth. Absence of sclerotized area. Pharyngeal armature without pigmentation, occupying the posterior fifth portion of the pharynx including dot-like teeth and lines. A light pigmentation is observed throughout the pharynx.

###### 3.3.1.1.2 Cervix (observed on paratype NEA0149.3)

Two cervical sensillae. Ventro-cervical sensilla not observable.

###### 3.3.1.1.3 Thorax

590 µm long. Brown sclerites. Mesonotum: presence of one post-alar seta; presence of three proepimeral setae; absence of the upper anepisiternal, lower anapisternal, anepimeral, metaepisternal, and metaepimeral setae; absence of setae on the anterior region of the katepisternum; absence of suture between metaepisternum and katepisternum. Metafurca mounted in lateral view on all specimens. Wings: length = 2056 μm, width = 661 μm, r5 = 1196 μm, α (r2) = 553 μm, β (r2+3) = 230 μm, δ = 87 μm, γ (r2+3+4) = 175 μm, ε (r3) = 684 μm, θ (r4) = 999 μm, π = 55 μm. Width/γ = 3.78. Anterior leg: procoxa = 345 µm; femur = 902 µm; tibia = 1082 µm; tarsomere I = 648 µm; sum of tarsomeres II-V= 699 µm.

###### 3.3.1.1.4 Abdomen

Tergites II-VI: presence of randomly distributed setae. Lack of setae on tergite VIII. Tergite IX without any protuberance. Cerci: 141 μm long. Setae not observed on sternite X.

###### 3.3.1.1.5 Genitalia

Spermathecal common duct 60 µm long with thicker walls roughly towards the basal 30 µm end. Opening in the genital atrium is enlarged (triangular shape). Common duct 6-7 µm wide at its middle and 22 µm wide at its basal opening. Spermathecal individuals ducts 59 μm long, 3-4 µm wide, with narrow thin wall exhibiting about 20 segments. Spermathecae: 41-46 µm long and about 12-13 µm wide. They are annealed with 15 rings. The terminal ring is larger and narrower (8-9 µm x 8-9 µm) than the other rings. Sessile head.

###### 3.3.1.2 Description of the male of *Ph. seowpohi* n. sp. (Fig 6)

The counts and measurements provided below are those of the paratype NEA0149.7. Measurements carried out on several specimens are indicated in S3 Table.

**Fig 6.**
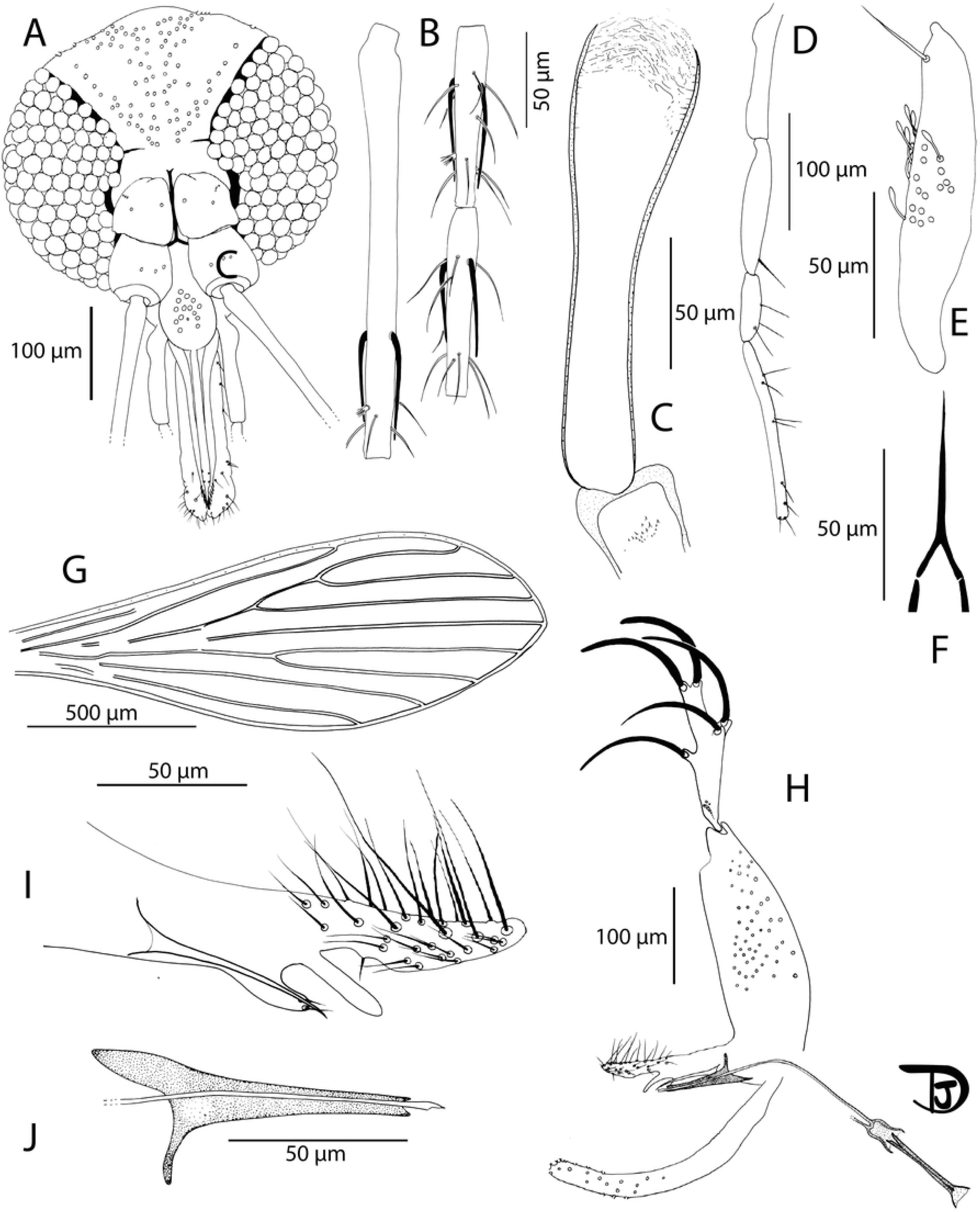
*Phlebotomus seowpohi* n. sp. male. A: head; B: flagellomeres 1, 2, and 3; C: pharynx and cibarium; D: palpi; E: 3rd palpal article; F: labial furca; G: wing; H: genitalia; I: trifurcated parameter and the accessory stick; J: parameral sheath and tip of the aedeagal ducts.

###### 3.3.1.2.1 Head

Occiput with two lines of well individualised setae. Clypeus: 122 µm long, 60 µm wide, with 13 big randomly distributed setae. Eyes: 245 µm long, 123 µm wide, with about 130 facets. Interocular sutures incomplete. Interantennal sutures do not reach the interocular sutures. Flagellomeres: f1 (=AIII) = 227 µm, f2 (=AIV) = 100 µm, f3 (=AV) = 100 µm, f12 (=AXIV) = 75 µm, f13 (=AXV) = 71 µm, f14 (=AXVI) = 80 µm. Flagellomere 1 longer than f2+f3. Presence of two ascoids from f1 to f8, and one from f9 to f13. Ascoidal formula: 2/f1-f18, 1/f9-f13. Long ascoids not reaching the next articulation. Ascoid of f2: 60.61 µm long. Length of this ascoid/length of f2 = 0.61. One distal papilla on f1 and f2. Absence of papillae on f3 to f8. Two papillae on f9, five on f10 and f11, six on f12 and f13, and five on f14. Several simple setae on all flagellomeres. Palps: p1 = 33 µm, p2 = 87 µm, p3 = 117 µm, p4 = 57 µm, p5 = 156 µm. Palpal formula: 1, 4, 2, 3, 5. Presence of a group of about 15 club-like Newstead’s sensilla in the middle of the third palpal segment. No Newstead’s sensilla on other palpal segments. Presence of one distal spiniform seta on p3, three on p4, and six on p5. Labrum: 178 µm long. f1/labrum = 1.28. Labial furca closed. Cibarium: presence of a few small teeth (denticles) in the centre. Absence of a sclerotized area. Pharyngeal armature occupying the posterior fifth portion of the pharnyx with small dot-like or line-like teeth arranged concentrically. Presence of a few lateral small teeth positioned anteriorly.

###### 3.3.1.2.2 Cervix

Two cervical sensillae on each side.

###### 3.3.1.2.3 Thorax

480 µm long. Brown sclerites. Mesonotum: presence of one post-alar seta; Presence of three proepimeral setae; absence of upper anepisiternal, lower anapisternal, anepimeral, metaepisternal, and metaepimeral setae; absence of setae on the anterior region of the katepisternum; absence of suture between metaepisternum and katepimeron. Metafurca with separated vertical arms. Wings: length = 1622 μm, width = 577 μm, r5 = 1009 μm, α (r2) = 448 μm, β (r2+3) = 195 μm, δ = 410 μm, γ (r2+3+4) = 145 μm, ε (r3) = 580 μm, θ (r4) = 836 μm, π = 67 μm. Width/γ = 3.97. Anterior leg: procoxa = 298 µm; femur = 740 µm; tibia = 1002 µm; tarsomere I = 635 µm; sum of tarsomeres ii-v = 650 µm. Median leg: mesocoxa = 329 µm; femur = 677 µm; tibia = 1164 µm; tarsomere I = 750 µm; sum of tarsomeres II-V = 663 µm. Posterior leg: metacoxa = 360 µm; femur = 772 µm; tibia = 1414 µm; tarsomere I = 831 µm; sum of tarsomeres II-V = 749 µm.

###### 3.3.1.2.4 Abdomen

Tergites II-VII: presence of randomly distributed setae. Presence of two tergal papillae (one per side) on tergites IV-VII.

###### 3.3.1.2.5 Genitalia

Absence of hypandrial apodeme (=abdominal rod).Gonocoxite: 258 µm long with a median tuft of about 30 internal setae. Absence of basal gonocoxal lobe. Gonostyle: 167 µm long with 5 thick spines (two terminal and three median ones).

Absence of accessory setae. Parameres: complex, 150 µm long, with a long upper lobe (70 µm long) exhibiting many setae, a brown shorter and narrow hairless intermediate lobe (30 µm long), and a small lower lobe (22 µm long) exhibiting two setae. Presence of a long (75 µm long), pale accessory spine between the paramere and the parameral sheath, narrow pointed at its tip. Parameral sheath: 87 µm long, blunt end. Aedeagal ducts: 235 µm long, smooth, beveled at the tip. Sperm pump: 126 µm long with a narrow ejaculatory apodeme 102 µm long. Epandrial lobes: 300 µm long, longer than the gonocoxite, without permanent setae. Cerci: 150 μm long.

#### 3.3.2 Description of *Sergentomyia gubleri* Depaquit, Vongphayloth, and Torno n. sp

Genus *Sergentomyia* França & Parrot.

We describe only the female. The male remains unknown.

##### Etymology

The epithet *gubleri* refers to Emeritus Professor Duane J. Gubler, Duke-National University of Singapore, in recognition of his significant contributions to public health entomology. His decades of work have enhanced the understanding and control of vector-borne diseases, and significantly improving public health outcomes worldwide.

Type locality: Dairy Farm (1°20’54.8”N 103°46’37.5”E), Singapore.

##### Type specimen

Female holotype (voucher NEA1119.7.BF1) will be deposited at the Laboratory of Entomology, The Natural History Museum, London, UK.

To meet the criteria of availability, the authors Depaquit, Vongphayloth, and Torno are responsible for the name *Sergentomyia gubleri* n. sp. and should be cited as the sole authority of this taxa, according to the Article 50 (1) of the lnternational Code of Zoological Nomenclature.

###### 3.3.2.1 Description of the female of *Se. gubleri* n. sp. (Fig 7)

Description and measurements indicated are those of the female holotype labelled NEA1119.7.BF1.

**Fig 7.**
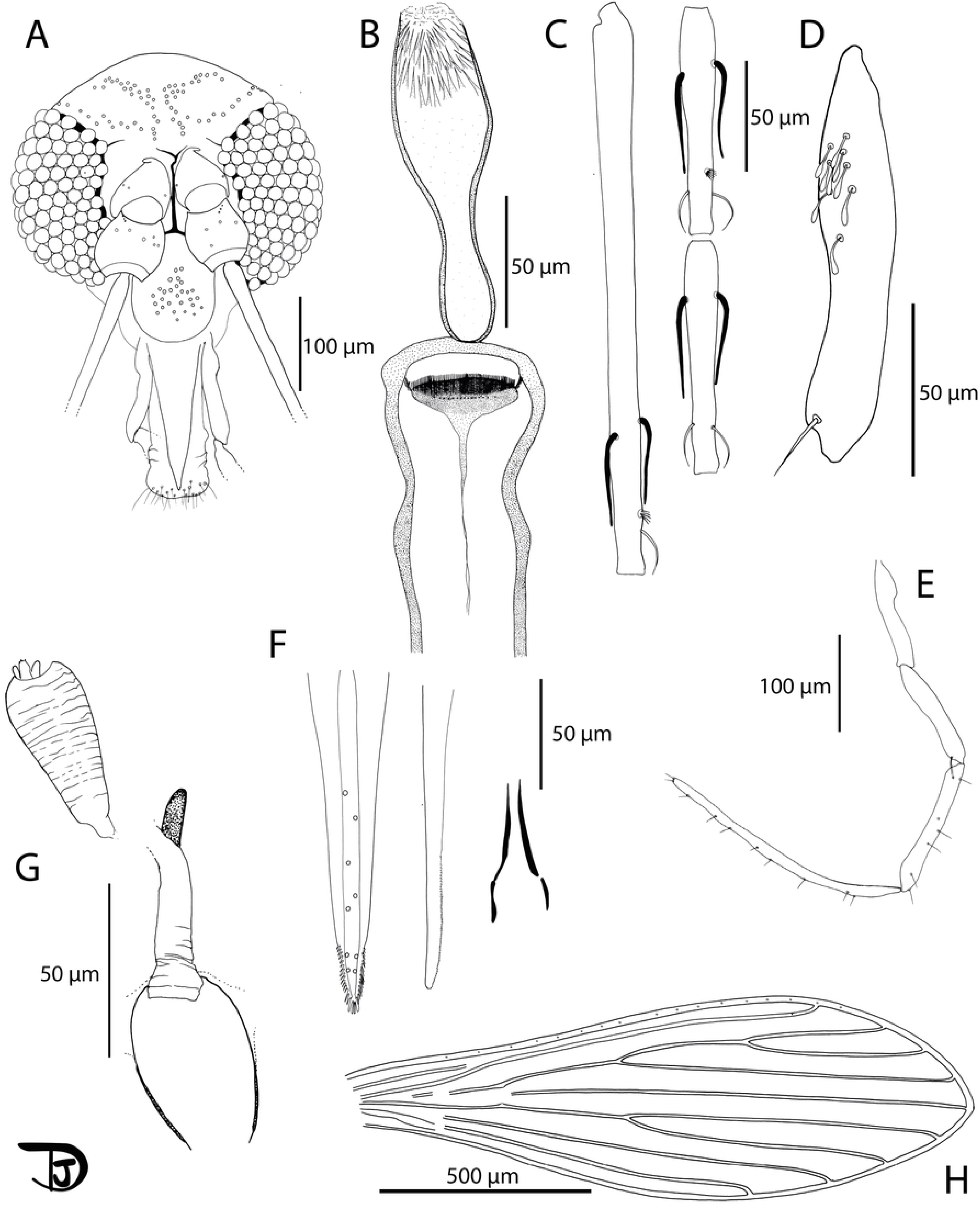
*Sergentomyia gubleri* n. sp. female. A: head; B: pharynx and cibarium; C: flagellomeres 1, 2, and 3; D: 3rd palpal article; E: palpi; F: mouth parts: labrum, mandible, and labial furca from left to right; G: genital furca and spermathecae; H: wing.

###### 3.3.2.1.1 Head

Occiput with two narrow lines of well individualised setae. Clypeus: 96 µm long84 µm wide, exhibiting 25 setae. Eyes: 177 µm long, 82 µm wide, with about 110 facets. Incomplete interocular sutures. Interantennal sutures do not reach the interocular ones. Flagellomeres: f1 (=AIII) = 250 µm, f2 (=AIV) = 101 µm, f3 (=AV) = 100 µm, f12 (=AXIV) = 72 µm, f13 (=AXV) = 55 µm, f14 (=AXVI) = 40 µm. Flagellomere 1 longer than f2+f3. Presence of 2 ascoids from f1 to f13. Ascoidal formula: 2/f1-f13. One distal papilla on f1, f2, f6, and f7. Absence of papillae on f3 to f5. Three papilla on f8 to f10 (two posterior and one median), four on f11 (two posterior, one median, and one distal), and five on f12 (two posterior, two median, and one distal). One simple seta on f1, two on f2 to f11, three on f12, and nine on f13. Palps: p1 = 45 µm, p2 = 67 µm, p3 = 113 µm, p4 = 139 µm, p5 = 257µm. Palpal formula: 1, 2, 3, 4, 5. Presence of a group of about eight club-like Newstead’s sensillae in the middle of the third palpal segment. No Newstead’s sensilla on other palpal segments. Presence of one apical spiniform seta on p3, seven on p4, and about ten on p5. Labrum: 142 µm long. f1/labrum = 1.76. Teeth of the maxillary lacinia not observable. Labial furca open. Cibarium armed with 68 posterior horizontal teeth and about 15 anterior vertical teeth. Presence of a distinctly dark oval sclerotized area, not reaching the posterior tip of the horizontal teeth. Well-developed pharyngeal armature occupying the posterior third part of the pharynx with about 40 lightly pigmented, long and pointed backwards oriented teeth.

###### 3.3.2.1.2 Cervix

Two cervical sensillae. Ventro-cervical sensilla not observable.

###### 3.3.2.1.3 Thorax

Brown sclerites. Mesonotum: absence of post-alar, proepimeral, upper anepisiternal, lower anapisternal, anepimeral, metaepisternal, and metaepimeral setae; absence of setae on the anterior region of the katepisternum. Metafurca with long separated vertical and horizontal arms. Wings: length = 1497 μm, width = 464 μm, r5 =1058 μm, α (r2) = 332 μm, β (r2+3) = 285 μm, δ = 176 μm, γ (r2+3+4) = 262 μm, ε (r3) = 431 μm, θ (r4) = 776 μm, π = 16 μm. Width/γ = 1.77. Anterior leg: procoxa = 255 µm; femur = 560 µm; tibia = 505 µm; tarsomere I = 260 µm; sum of tarsomeres II-V = 452 µm. Median leg: mesocoxa = 262 µm; femur = 607 µm; tibia = 626 µm; tarsomere I = 290 µm; sum of tarsomeres II-V = 489 µm. Posterior leg: metacoxa = 265 µm; femur = 607 µm; tibia = 788 µm; tarsomere I = 355 µm; sum of tarsomeres II-V = 727 µm.

###### 3.3.2.1.4 Abdomen

Tergites II-VI: presence of randomly distributed setae. Lack of setae on tergite VIII.

Tergite IX without any protuberance. Cerci: 149 μm long. Setae not observed on sternite X.

###### 3.3.2.1.5 Genitalia

Spermathecae: 52 µm long and about 26 µm wide with wrinkled thin walls. Wider at its apex than at the proximal part. Sessile head.

#### 3.3.3 Description of *Sergentomyia leechingae* Depaquit, Vongphayloth, and Ding n. sp

Genus *Sergentomyia* França & Parrot.

##### Etymology

The epithet *leechingae* refers to Associate Professor Lee-Ching Ng, Group Director of the Environmental Health Institute, NEA, Singapore, in recognition of her significant contributions to Singapore’s public health and her impactful work in the prevention and control of vector-borne diseases both in Singapore and globally.

Type locality: Lim Chu Kang, Singapore (1°26’12.6”N 103°42’50.6”E)

##### Type specimen

female holotype (voucher NEA0118.1) and one male paratype (voucher NEA0118.4) will be deposited at the Laboratory of Entomology, The Natural History Museum, London, UK.

One female paratype (voucher NEA0299) will be deposited at the Environmental Health Institute, National Environment Agency, Singapore.

To meet the criteria of availability, the authors Depaquit, Vongphayloth, and Ding are responsible for the name *Sergentomyia leechingae* n. sp. and should be cited as the sole authority of this taxa, according to the Article 50 (1) of the lnternational Code of Zoological Nomenclature.

###### 3.3.3.1 Description of the female of *Se. leechingae* n. sp. (Fig 8)

Description and measurements indicated are those of the female holotype labelled NEA0118.1.

**Fig 8.**
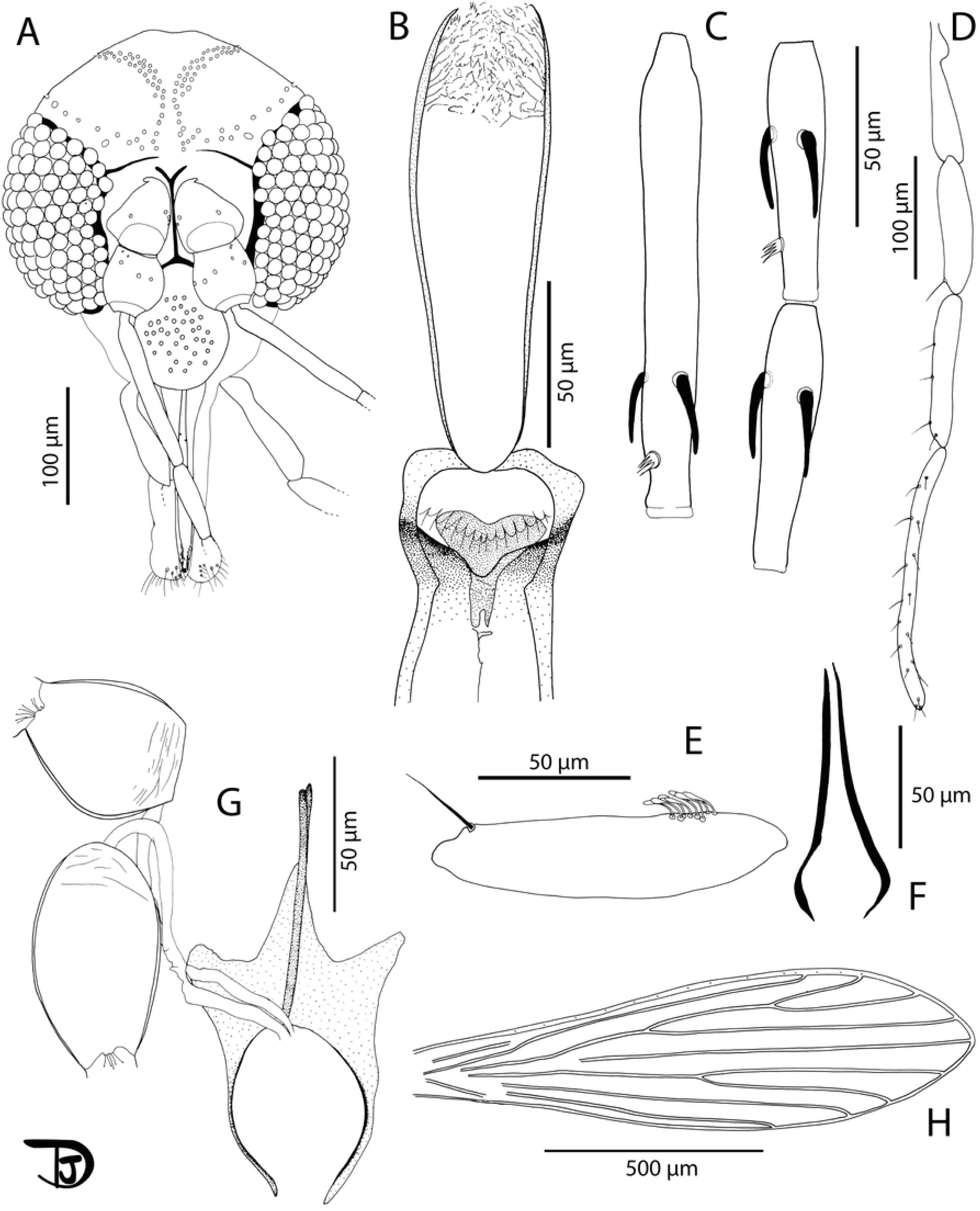
*Sergentomyia leechingae* n. sp. female. A: head; B: pharynx and cibarium; C: flagellomeres 1, 2, and 3; D: palpi; E: 3rd palpal article; F: labial furca; G: genital furca and spermathecae; H: wing.

###### 3.3.3.1.1 Head

Occiput with two narrow lines of well individualised setae. Clypeus: 118 µm long, 70 µm wide, exhibiting 39 setae. Eyes: 175 µm long, 76 µm wide, with about 100 facets. Incomplete interocular sutures. Interantennal sutures do not reachthe interocular sutures. Flagellomeres: f1 (=AIII) = 125 µm, f2 (=AIV) = 70 µm, f3 (=AV) =65 µm, f12 (=AXIV) = 45 µm, f13 (=AXV) = 39 µm, f14 (=AXVI) = 38 µm. Flagellomere 1 shorter than f2+f3. Presence of two short ascoids from f1 to f13, not reaching the next articulation. Ascoidal formula: 2/f1-f13. One distal papilla on f1 and f2. Absence of papillae on f3 to f12. Three papillae on f12 and f13, and two on f14. Palps: p1 = 41 µm, p2 = 65 µm, p3 = 98 µm, p4 = 120 µm, p5 = 206 µm. Palpal formula: 1, 2, 3, 4, 5. Presence of a group of less than 10 club-like Newstead’s sensilla in the proximal part of p3. No Newstead’s sensilla on other palpal segments. Presence of one distal spiniform setae on p3, five on p4, and about 15 on p5.Labrum: 147 µm long. f1/labrum = 0.85. Mouth parts not observable. Labial furca open. Cibarium armed with 14 posterior teeth pointed backwards. Absence of anterior teeth. Presence of sclerotized area, heart-like with a bifid anterior expansion. Presence of a notch in the middle of the cibarium, just below the area where anterior teeth would have been located. Inconspicuous pharyngeal armature with some pointed teeth oriented backwards and some tiny teeth arranged along numerous lines. Armature occupies the posterior quarter of the pharynx.

###### 3.3.3.1.2 Cervix

Two cervical sensillae on each side. Ventro-cervical sensilla not observed.

###### 3.3.3.1.3 Thorax

525 µm long. Light brown sclerites. Mesonotum: absence of post-alar, proepimeral, upper anepisiternal, lower anapisternal, anepimeral, metaepisternal, and metaepimeral setae; setae on the anterior region of the katepisternum not observed. Metafurca mounted in lateral view on all specimens. Wings: length = 1122 μm, width = 369 μm, r5 = 887 μm, α (r2) = 160 μm, β (r2+3) = 268 μm, δ = 39 μm, γ (r2+3+4) = 220 μm, ε (r3) = 269 μm, θ (r4) = 627 μm, π = 82 μm. Width/γ = 1.68. Anterior leg: procoxa = 235 µm; femur = 503 µm; tibia = 430 µm; tarsomere I = 220 µm; sum of tarsomeres II-V = 419 µm.

###### 3.3.3.1.4 Abdomen

Tergites II-VI: presence of randomly distributed setae. Lack of setae on tergite VIII. Tergite IX without any protuberance. Cerci: 90 μm long. Setae not observed on sternite X.

###### 3.3.3.1.5 Genitalia

Spermathecal common duct absent. Spermathecal individual ducts 125 µm long. Spermathecae: 57 µm long and 33 µm wide, oblong or oval with slight streaks at the base. Sessile head.

###### 3.3.3.2 Description of the male of *Se. leechingae* n. sp. (Fig 9)

The counts and measurements provided below are those of the paratype NEA0118.4.

**Fig 9.**
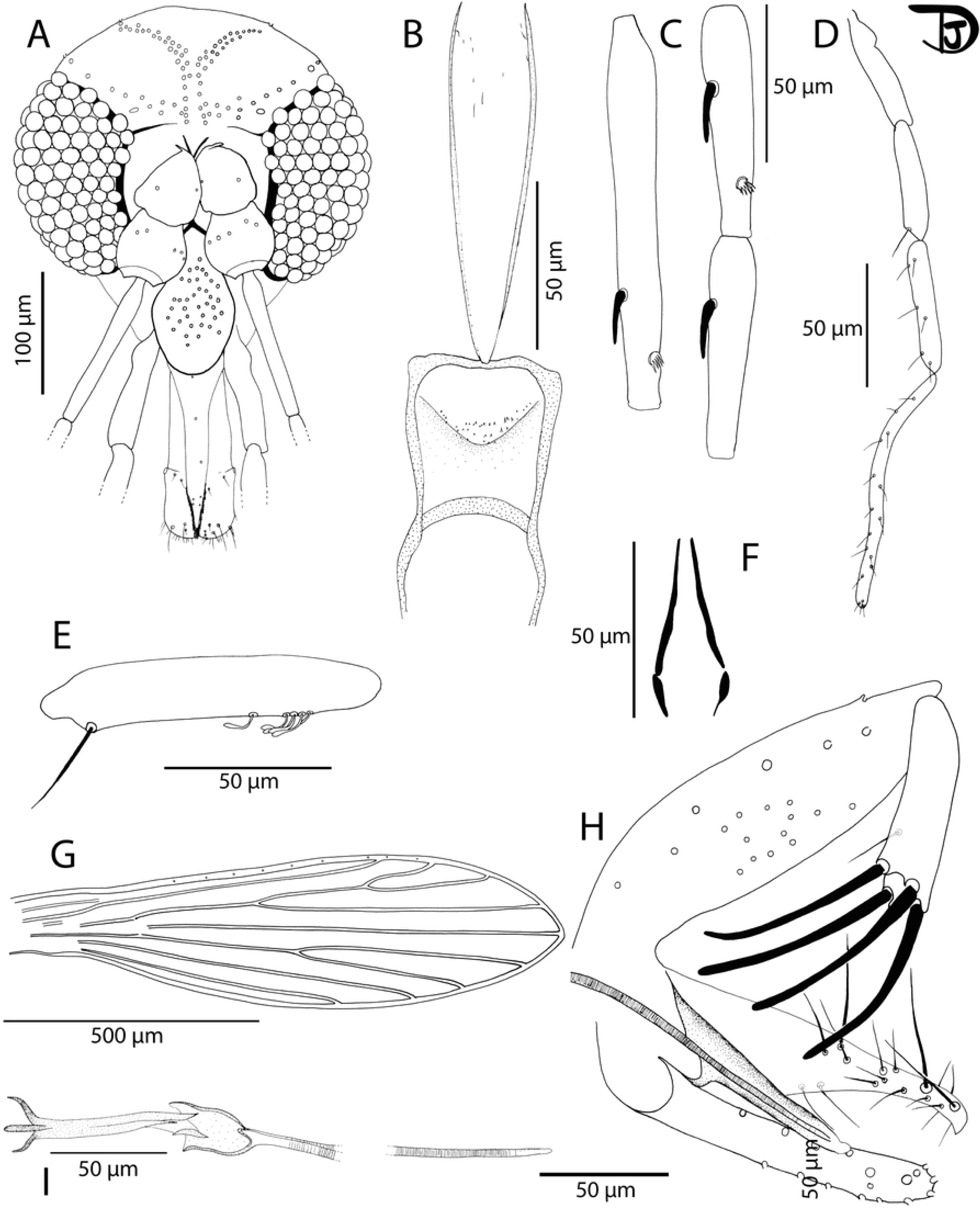
*Sergentomyia leechingae* n. sp. male. A: head; B: pharynx and cibarium; C: flagellomeres 1, 2, and 3; D: palpi; E: 3rd palpal article; F: labial furca; G: wing; H: genitalia; I: sperm pump and tip of the aedeagal ducts.

###### 3.3.3.2.1 Head

Occiput with two lines of well individualised setae. Clypeus: 117 µm long, 63 µm wide, with 34 big randomly distributed setae. Eyes: 165 µm long, 82 µm wide, with about 90 facets. Interocular sutures incomplete, not observable. Interantennal sutures do not reach the interocular sutures. Flagellomeres: f1 (=AIII) = 116 µm, f2 (=AIV) = 69 µm, f3 (=AV) = 68 µm. Flagellomere 1 shorter than f2+f3. Presence of one short ascoid from f1 to f7, not reaching the next articulation. Antenna is broken after f7. One distal papilla on f1, f2. Absence of papilla on f3 to f8. Palps: p1 = 33 µm, p2 = 87 µm, p3 = 117 µm, p4 = 57 µm, p5 = 156 µm. Palpal formula: 1, 4, 2, 3, 5. Presence of six club-like Newstead’s sensillae at the proximal third of the palpal segment. No Newstead’s sensilla on other palpal segments. Presence of one distal spiniform seta on p3, six on p4, and about 15 on p5. Labrum: 129 µm long. f1/labrum = 0.89. Labial furca open. Cibarium: presence of a few small randomly arranged teeth (denticles) in the middle. Absence of a sclerotized area. Very few teeth observed at the posterior part of the pharynx.

###### 3.3.3.2.2 Cervix

Two cervical sensillae on each side. Ventro-cervical sensilla not observed

###### 3.3.3.2.3 Thorax

Brown sclerites. Mesonotum: absence of post-alar, proepimeral, upper anepisiternal, lower anapisternal, anepimeral, metaepisternal, and metaepimeral setae; absence of setae on the anterior region of the katepisternum. Metafurca with separated vertical arms. Wings: length = 1030 μm, width = 292 μm, r5 = 777 μm, α (r2) = 139 μm, β (r2+3) = 206 μm, δ = 37 μm, γ (r2+3+4) = 178 μm, ε (r3) = 233 μm, θ (r4) = 533 μm, π = 74 μm. Width/γ = 1.64. Legs broken.

###### 3.3.3.2.4 Abdomen

Tergite II-VII: presence of randomly distributed setae. Presence of two tergal papillae (one per side) on tergites IV-VII.

###### 3.3.3.2.5 Genitalia

Gonocoxite: 173 µm long with around ten internal setae scattered randomly around the medial region. Absence of basal gonocoxal lobe. Gonostyle: 75 µm long with 4 thick terminal spines. Presence of an accessory seta on the distal quarter of the gonostyle. Parameres: simple, 127 µm long. Parameral sheath: 89 µm long. Aedeagal ducts: 240 µm long. Tip of the ducts rounded. Sperm pump: 96 µm long with a narrow ejaculatory apodeme 82 µm long. Epandrial lobes: 138 µm long, shorter than the gonocoxite. Cerci: 150 μm long.

#### 3.3.4 Description of *Seregentomyia retrocalcarae* Depaquit, Vongphayloth, and Torno n. sp

Genus *Sergentomyia* França & Parrot.

We describe only the female. The male remains unknown.

##### Etymology

From Latin *retro*, oriented backwards, and *calcar*, spur, referring to spurs on the ascoids which are oriented backwards. Type locality: Lim Chu Kang (1°26’35.7”N 103°42’28.6”E), Singapore.

##### Type specimen

Female holotype (voucher NEA0120.1) will be deposited at the laboratory of Entomology, The Natural History Museum, London, UK. One female paratype (voucher NEA0008) will be deposited at the Environmental Health Institute, National Environment Agency, Singapore.

To meet the criteria of availability, the authors Depaquit, Vongphayloth, and Torno are responsible for the name *Sergentomyia retrocalcarae* n. sp. and should be cited as the sole authority of this taxa, according to the Article 50 (1) of the lnternational Code of Zoological Nomenclature.

###### 3.3.4.1 Description of the female of *Se. retrocalcarae* n. sp. (Fig 10)

Description and measurements indicated are those of the female holotype labelled NEA0120.1 except those of the genitalia observed on paratype NEA0008.

**Fig 10.**
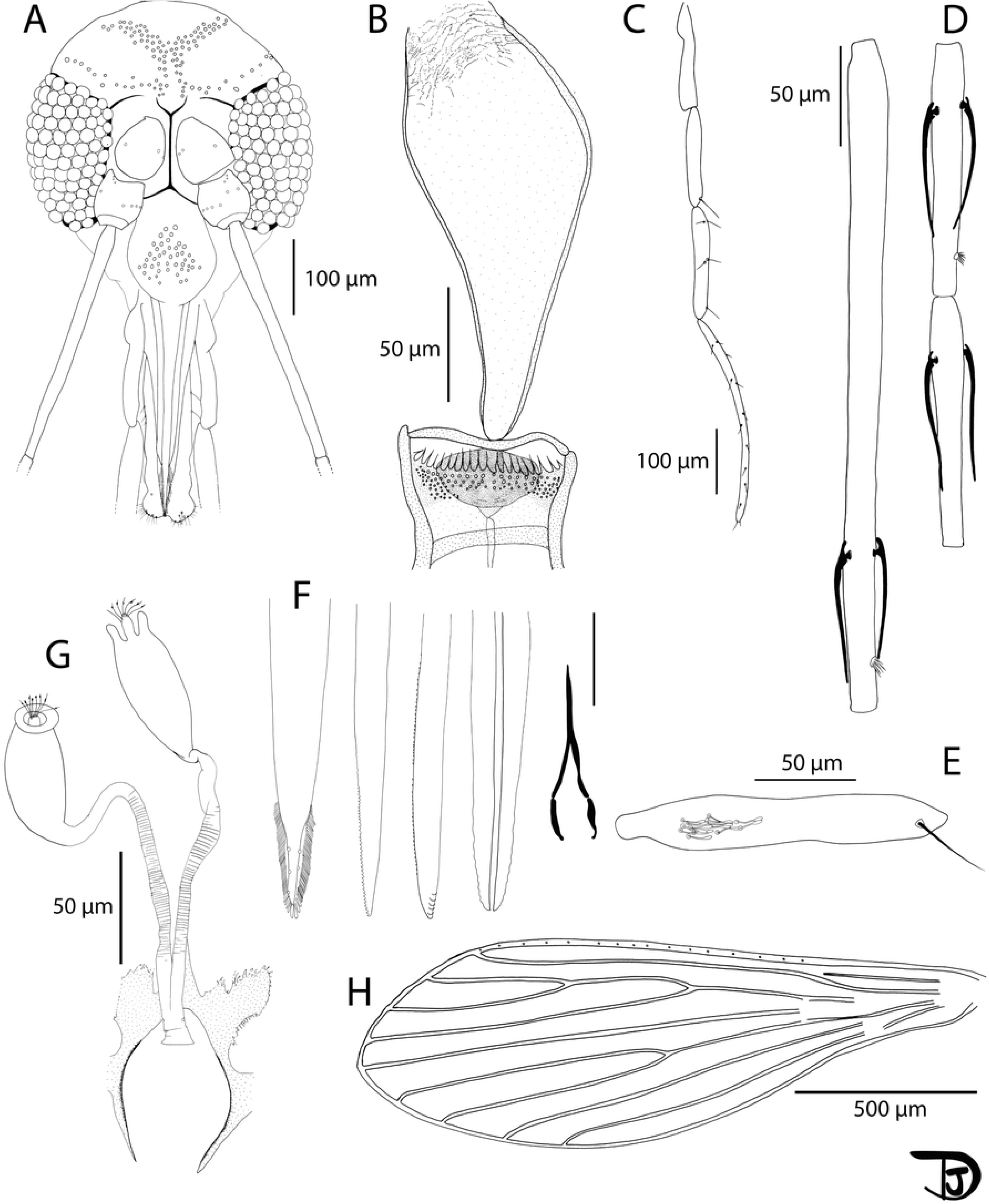
*Sergentomyia retrocalcarae* n. sp. female. A: head; B: pharynx and cibarium; C: palpi; D: flagellomeres 1, 2, and 3; E: 3rd palpal article; F: mouth parts: labrum, mandible, lacinia, hypopharynx and labial furca from left to right; G: genital furca and spermathecae; H: wing.

###### 3.3.4.1.1 Head

Occiput with two narrow lines of well individualised setae. Clypeus: 138 µm long, 65 µm wide, exhibiting 44 setae. Eyes: 205 µm long, 98 µm wide, with about 80 facets. Interocular sutures incomplete. Interantennal sutures do not reach the interocular sutures. Flagellomeres: f1 (=AIII) = 319 µm, f2 (=AIV) = 120 µm, f3 (=AV) = 119 µm, f12 (=AXIV) = 75 µm, f13 (=AXV) = 58 µm, f14 (=AXVI) = 53 µm. Flagellomere 1 longer than f2+f3. Presence of two ascoids from f1 to f13. Ascoidal formula: 2/f1-f13. One apical papilla on f1 and f2. Absence of papilla on f3 to f5. One papilla on f6 to f11, and four on f12 to f14. Palps (observed on NEA0008): p1 = 57 µm, p2 = 104 µm, p3 = 149 µm, p4 = 174 µm, p5 = 306 µm. Palpal formula: 1, 2, 3, 4, 5. Presence of a dozen club-like Newstead’s sensillae located at the proximal third of the third palpal segment. No Newstead’s sensilla on other palpal segments. Presence of one apical spiniform seta on p3, five on p4, and thirteen on p5. Labrum: 258 µm long. f1/labrum = 1.24. More than 50 lateral teeth on each side of the labrum. Mandible exhibiting about 40 lateral teeth. Hypopharynx exhibiting about ten waved teeth on each side. Maxillary lacinia exhibiting five external teeth and about 50 internal teeth. Labial furca open. Cibarium armed with 19 horizontal posterior horizontal teeth pointed backwards. Presence of many anterior teeth. Presence of an oval sclerotized area does not reach the lateral walls of the cibarium but reaching the tip of the horizontal teeth. Pharynx slightly pigmented under the armature. Pharyngeal armature at the posterior fifth of the pharynx, consisting of minute teeth forming concentric curves.

###### 3.3.4.1.2 Cervix

Two cervical sensillae. Ventro-cervical sensilla not observable.

###### 3.3.4.1.3 Thorax

Brown sclerites. Mesonotum: absence of post-alar, proepimeral, upper anepisiternal, lower anapisternal, anepimeral, metaepisternal, and metaepimeral setae; absence of setae on the anterior region of the katepisternum. Metafurca with long separated vertical arms. Wings: length = 1600 μm; width = 554 μm, r5 = 1196 μm, α (r2) = 348 μm, β (r2+3) = 329 μm, δ = 145 μm, γ (r2+3+4) = 317 μm, ε (r3) = 465 μm, θ (r4) = 860 μm, π = 87 μm. Width/γ=1.74. Anterior leg: procoxa = 148 µm; femur = 338 µm; tibia = 334 µm; and tarsomere I = 159 µm; sum of tarsomeres II-V = 258 µm. Median leg: mesocoxa = 232 µm; femur = 666 µm; tibia = 837 µm; and tarsomere I = 369 µm; sum of tarsomeres II-V = 543 µm. Posterior leg: metacoxa = 285 µm; femur = 728 µm; tibia =1064 µm; and tarsomere I = 491 µm; sum of tarsomeres II-V = 613 µm.

###### 3.3.4.1.4 Abdomen

Tergites II-VI: presence of randomly distributed setae. Six setae on tergite VIII. Tergite IX without any protuberance. Cerci: 126 μm long. Setae not observed on sternite X.

###### 3.3.4.1.5 Genitalia

Spermathecal common duct short, 10 µm long. Spermathecal individual ducts 145 µm long. Spermathecae (measurement taken on NEA0008): 57 µm long and about 33 µm wide, oblong and smooth. Sessile head.

## 4. Discussion

This study reports the first record of eight species of *Phlebotomus* and *Sergentomyia* sand flies in Singapore. They include *Ph. stantoni*, *Se. barraudi* group, *Se. iyengari* group, *Se. whartoni*, and the discovery of four new species – *Ph. seowpohi* n. sp., *Se. gubleri* n. sp., *Se. leechingae* n. sp., and *Se. retrocalcarae* n. sp. The new species are described through the integrative taxonomic approach using both morphological and molecular diagnostic methods.

Majority of sand flies collected are assigned to the new species, *Phlebotomus (Euphlebotomus) seowpohi* n. sp.. According to previous reviews (Seccombe *et al.* (1993) [62], Leng & Lewis (1987) [63], Javadian et al. (1997) [64], Muller *et al.* (2007) [65], and Depaquit *et al*. (2009) [66]), the subgenus *Euphlebotomus* includes the following species: *Ph. argentipes s. l.*; *Ph. autumnalis*; *Ph. barguesae*; *Ph. caudatus*; *Ph. kiangsuensis*; *Ph. mascomai*; *Ph. mesghalii*; *Ph. nadimi*; *Ph. philippinensis*; *Ph. gouldi* stat. nov.; *Ph. tumenensis;* and *Ph. yunshengensis*.

We did not consider *Ph. lengi* originally classified in the subgenus *Larroussius* (Zhang *et al.*, 1994) as a *Euphlebotomus* species [67]. This was previously proposed by Zhang & Leng (1997) [68], but additional examination of both male and female specimens using morphological and molecular approaches suggested otherwise [66]. Furthermore, *Phlebotomus argentipes s.l.* could be a complex of three species including *Ph. argentipes s. st., Ph. annandalei*, and *Ph. glaucus* [69–72].

The goal of the present study is not to focus on the validity of these species as their statuses are not adopted by all authors. Instead, we propose to recognize *Ph. seowpohi* as a member of the subgenus *Euphlebotomus.* The ratio between the length of its ascoid on the second flagellomere and the length of its second flagellomere (asc/f2 = asc/AIV) was investigated primarily as this character is considered as the most discriminant by authors working on the variability within the *Ph. argentipes* complex. Female *Ph. seowpohi* n. sp. specimens have asc/f2 ratio of 0.71, while males have a ratio of 0.61. According to literature, *Ph. argentipes* and *Ph. annandalei* are species with a short f2 ascoid (asc/f2 < 0.5), while *Ph. glaucus* has long ascoids (asc/f2 ranges between 0.50 and 0.60) [71,72]. Attempts were made to check this character in the type specimens. Unfortunately, it was not possible to have access to *Ph. argentipes* nor *Ph. glaucus* type specimen. Additionally, no type specimen was previously deposited for *Ph. annandalei*.

*Ph. seowpohi* n. sp. includes two distinct morphological characters. Firstly, both sexes do not exhibit any papilla on the 3^rd^ flagellomere. This is completely unusual within the genus *Phlebotomus*. Although old descriptions did not mention the absence nor presence of such sensilla on the 3^rd^ flagellomere, recent observations carried out on the *Phlebotomus* and *Sergentomyia* genera highlighted the presence of such sensilla for the genus *Phlebotomus* and its absence for the genus *Sergentomyia* [13,31,65,66,73–75]. This unusual observation suggests that *Ph. seowpohi* n. sp. shares a character previously known for the genus *Sergentomyia*. Secondly, both sexes exhibit one post-alar setae on the mesonotum. This presence seems unusual but has rarely been documented in taxonomic descriptions published before the experts’ guidelines [54].

To aid future researchers in the identification of *Ph. seowpohi* n. sp. amidst taxonomic challenges in the subgenus *Euphlebotomus*, the differential diagnoses of *Ph. seowpohi* n. sp. from other members of the subgenus are indicated in the following paragraphs.

Regarding the females, the pharyngeal armature of *Ph. seowpohi* n. sp. is colourless whereas it is pigmented in the following *Euphlebotomus* species: *Ph. barguesae*, *Ph. gouldi*, *Ph. kiangsuensis*, *Ph. mascomai*, and *Ph. philippinensis*. The 3^rd^ flagellomere of *Ph. mesghalii* (170-180 µm long) is shorter than those of *Ph. seowpohi* n. sp. (>220 µm long). The spermathecal ducts of *Ph. seowpohi* n. sp. are smooth whereas those of *Ph. yunshengensis* are tapeworm-like [76]. The spermathecal individual ducts of *Ph. autumnalis* are longer than the common duct whereas those of *Ph. seowpohi* n. sp. are shorter than the common duct. The spermatheca of *Ph. tumenensis* is wide and bulbous with faint striations near duct, but without distinct rings. To our knowledge, female specimens of *Ph. caudatus* and *Ph. nadimi* were never described.

Regarding the males, *Ph. autumnalis*, *Ph. caudatus*, *Ph.* mesghalii, and *Ph. nadimi* have a short main paramere lobe. No accessory spine between the parameral sheath and the paramere has been reported in the description of *Ph. mascomai* and the antennal formula of these species are different [65]. The genitalia of *Ph. tumenensis* does not exhibit any accessory spine between the parameral sheath and the paramere. *Ph. yunshengensis* exhibits an important tuft of setae on the gonocoxite. The males of *Ph. barguesae* have a blunt-end top of their parameral sheath not observed in the subgenus *Euphlebotomus.* The ratio between the length of the 3^rd^ flagellomere and the length of the labrum (f3/labrum) differentiate *Ph. seowpohi* n. sp. (about 1) from *Ph. philippinensis* and *Ph. gouldi* (>1.7). The thickness of the middle lobe of the paramere of *Ph. kiangsuensis* should differentiate it from *Ph. seowpohi* n. sp. according to Lewis (1982) but the taxonomic status of this species needs further clarification [48].

*Ph. seowpohi* n. sp. differs from *Ph. argentipes s. st.* by the relative length of their ascoids. The ascoids are long in *Ph. seowpohi* n. sp. and short for *Ph. argentipes*. The differential diagnosis with closely related species of the complex remains impossible until further taxonomic revision of the species complex. The best way to differentiate *Ph. seowpohi* n. sp. from *Ph. argentipes s. l.* will be through their highly divergent *cytochrome b* DNA sequences [77–79]. Molecular analyses on the *cytb* and *COI* gene fragment also suggests the validity of *Phlebotomus seowpohi* n. sp. as a new species. Sequences of males and females showed high homology with little variation between sequences. Pairwise intraspecific p-distances were 0.001 and 0.001 for *cytb* and *COI* sequences, respectively (Table 2 and S1 Table). This, together with the sequences forming monophyletic clades with high branch support (Fig 3 and S2 Fig), provides strong evidence that the sequences represent the same species and is distinctly unique from other genetically related species. Moreover, pairwise interspecific p-distances between sequences of *Ph. seowpohi* n. sp. and members of *Ph. (Euphlebotomus)* spp. had a minimum value of 0.125 and 0.146 for *cytb* and *COI* respectively (Table 2 and S1 Table). This suggests that the genetic sequences of *Ph. seowpohi* n. sp. is at least 12.5% different when compared with sequences of species from the same subgenus. This value is greater than the generally accepted DNA barcoding gap of about 3% used to discriminate sand fly species [77,80,81]. Hence, *cytb* and *COI* sequences generated in this study and deposited on Genbank will be an asset for entomological biosurveillance and molecular identification. Considering that molecular DNA barcoding requires much less technical expertise than species identification through morphological identification [81], future researchers can match their sequences to those deposited online for rapid species identification.

During the review of specimen inventories, it was determined that *Ph. seowpohi* n. sp. was also recorded in Laos. Specimens from Laos matched the specimens from Singapore in both morphological and molecular data (Fig 3 and S2 Figure). The distribution range of this new species thus requires further evaluation. *Phlebotomus seowpohi* n. sp. could potentially be widely distributed across SEA. In light of the description of *Ph. seowpohi* n. sp. in both countries, previous records of *Ph. argentipes* s. l. in SEA therefore remain doubtful. We propose a systematic taxonomic study of all the specimens caught in this region in order to ascertain the distribution of these species and to detect possible *Ph. argentipes* misidentifications in previous entomological surveys.

In Bangladesh, India, Nepal, and Bhutan, *Ph. argentipes* is an important vector of *Leishmania donovani*, and is recognized as the primary vector responsible for transmitting human visceral leishmaniasis in the South Asian region [82,83]. The discovery of *Ph. seowpohi* n. sp. has important public health significance, as findings from our integrative taxonomy analyses suggest that *Ph. seowpohi* may be closely related to *Ph. argentipes*. It is probable that *Ph. seowpohi* may serve as the vector of *Leishmania donovani* in SEA in the absence of the primary vector, *Ph. argentipes*. Comparative studies between *Ph. seowpohi* n. sp. and its *Leishmania*-transmitting relative can provide insights into the genetic and environmental factors that influence vector competence. Hence, further studies on *Ph. seowpohi* n. sp. are imperative to reveal whether it poses a similar threat as a vector of leishmaniasis.

Besides *Ph. seowpohi* n. sp., *Ph. stantoni* was the only other *Phlebotomus* species collected in Singapore. This species is known to be widespread across SEA [24,25,30,84]. Based on the maximum-likelihood tree generated from this study, the specimens from Singapore clearly clustered together with other *Ph. stantoni* collected from the region.

Unlike the genus *Phlebotomus,* which is well characterized, the genus *Sergentomyia* is considered a catch-all group by many phlebotomine sand fly taxonomists [8]. Its systematics remain debatable and is in need of revision. It is for this reason that we do not propose the inclusion of the new species we described into any subgenus. Useful taxonomic notes on *Sergentomyia* species collected in Singapore, including the newly described species, can be found in the following paragraphs.

*Sergentomyia gubleri* n. sp. exhibits many cibarial teeth similar to *Se. rudnicki* and *Se. brevicaulis*, which are members of the *Se. barraudi* group. However, this new species is molecularly distant from all other members of the group. It can be distinguished by the length of the first flagellomere as well as by the number of teeth constituting the comb-like cibarial armature (Table 4 and Fig 11).

**Fig 11.**
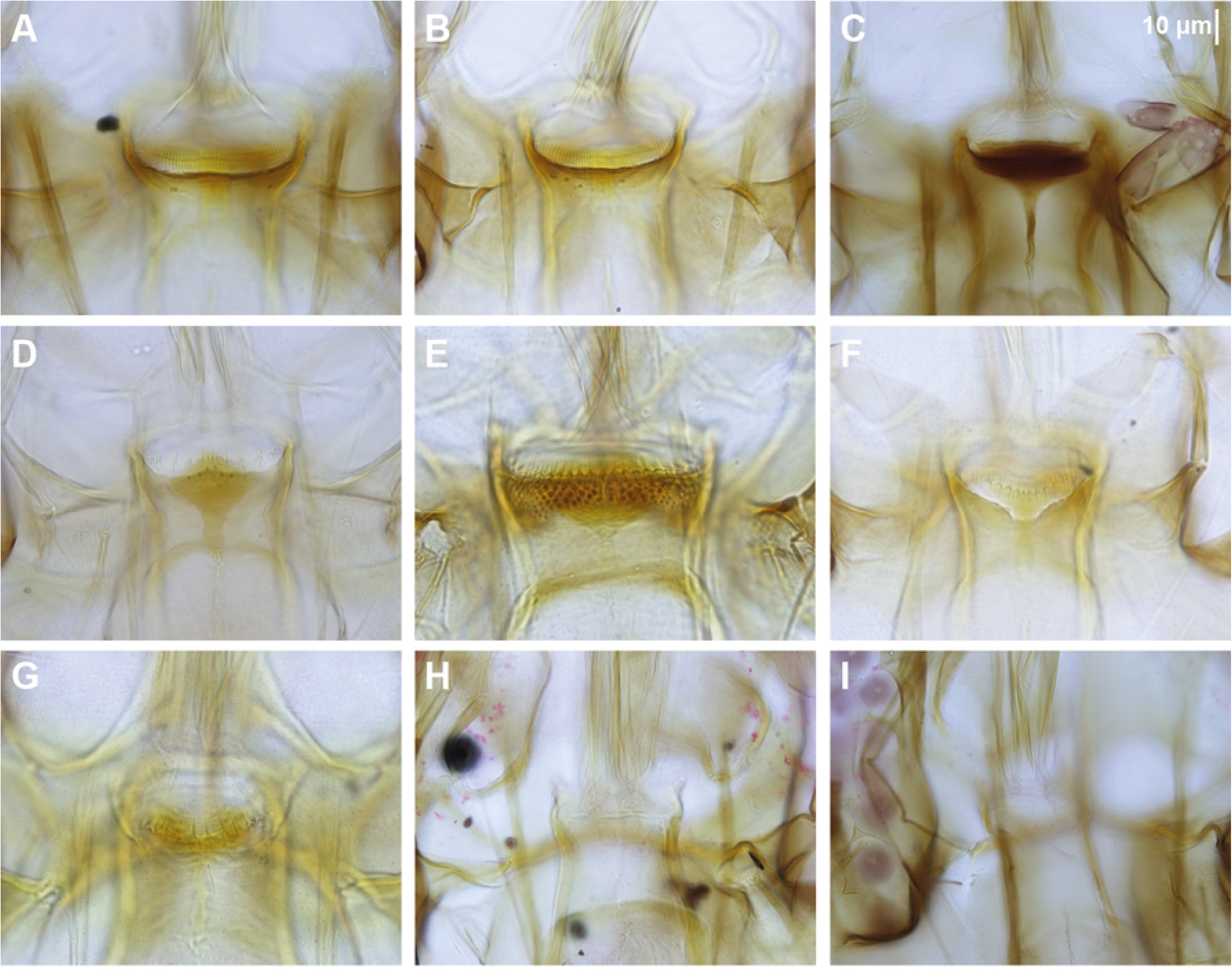
Cibarium of female sand flies collected in Singapore. All photos are at the same scale. A: *Sergentomyia barraudi* group NEA0501.1; B: *Se. barraudi* group NEA0009.2; C: *Se. gubleri* n. sp. NEA1119.1.BF7; D: *Se. iyengari* group NEA0283; E: *Se. retrocalcarae* n. sp. NEA0120.1; F: *Se. leechingae* n. sp. NEA0118.1; G: *Se. whartoni* NEA0069.1; H: *Phlebotomus stantoni* NEA0569; I: *Ph. seowpohi* n. sp. NEA0149.3.

**Table 4.**
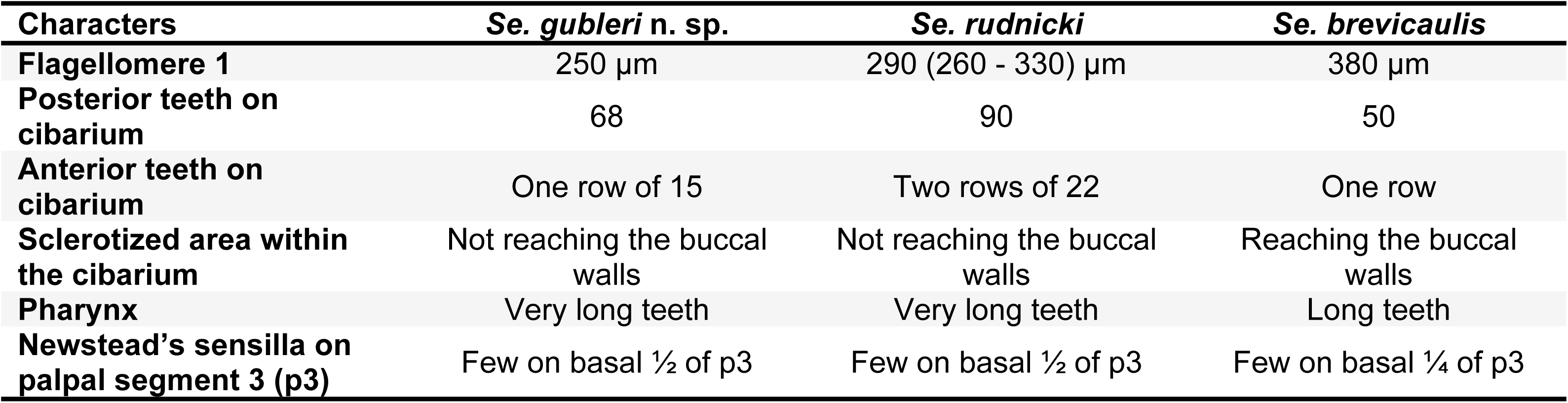
Differential diagnostic characteristics of *Se. gubleri* n. sp. compared against closely related species *Se. rudnicki* and *Se. brevicaulis*.

*Sergentomyia leechingae* n. sp. exhibits a cibarial notch similar to *Se. babu*, *Se. insularis*, *Se. baghdadis*, and *Se. shortii*, but is less developed (Fig 11). These species can be distinguished by shape of their cibarial notch together with the number of teeth on the cibarial armature as indicated in Table 5.

**Table 5.** Differential characters species. of female Se. leechingae n. sp. and related.

| Characters | *Se. leechingae*<br>n. sp | *Se. babu* | *Se. insularis* | *Se. baghdadis* | *Se. shortii* |
| --- | --- | --- | --- | --- | --- |
| Posterior teeth on cibarium | 14 | 24 - 34 | 45 - 50 | 16 - 20 | 10 - 14 on a straight line |
| Cibarial notch | Shallow | Deep | Deep | Deep | Shallow |

*Sergentomyia retrocalcarae* n. sp. exhibits spurs on the ascoids that are oriented backwards on the flagellomeres. It also has a cibarium which has many anterior teeth. However, the number of cibarial teeth for this species is less than those of *Se. gombaki* and *Se. arboris,* both of which also exhibit spurs on the ascoids. The differential diagnosis to closely related species is provided in Table 6.

**Table 6.** Differential characters of female *Se. retrocalcarae* n. sp. and related species.

| Characters | *Se. retrocalcarae*<br>n. sp. | *Se. gemmea* | *Se. malayae* | *Se raynali* |
| --- | --- | --- | --- | --- |
| Ascoid | Short spur | Short spur | Very long spur | Short spur |
| Anterior teeth on cibarium | Three rows, first row not very large | Three rows, first row very large | Three rows, first row not very large | Two rows, first row not very large. Few anterior teeth. |

The *Sergentomyia barraudi* group is undeniably a species complex which should be explored further by investigating both morphological and molecular characters of individuals collected from various populations across its distribution area. Morphological variability can be observed from the varying number and distribution of cibarial teeth (Fig 11), while the molecular variability is demonstrated by genetic differences between its mitochondrial barcodes (Fig 4, S2 Fig, Table 3, and S2 Table). The specimens from Singapore have more posterior teeth (56-60) than those of the original description (40) [85]. Hence, we identified the specimen collected in the current study as belonging to *Se. barraudi* group, as a more precise identification could be erroneous.

Many records of *Se. iyengari* have been made in South-Eastern Asia. However, this species was described from the southernmost part of India, which is not near the SEA region [86]. In our opinion, all records of *Se. iyengari* in SEA should in fact refer to either *Se. khawi* or *Se. hivernus*. The species *Se. tambori* Lewis & Jeffery, 1978 described from Malaysia could also be included in the *iyengari* group. As with the *barraudi* group discussed above, there is a need to revise the *iyengari* group by including various populations from India and Southeast Asia. In order to avoid further confusion and taxonomic errors, we prefer to identify specimens collected from Singapore as belonging to the *Se. iyengari* group. A photo of the cibarial armature of the specimen collected in this study is provided in Fig 11.

The specimen labelled NEA0069.1 resembles *Se. whartoni* described from continental Malaysia, about 200 km north of Singapore [87]. Its morphological characters match the original description of *Se. whartoni*. Even though we observed that the *COI* barcode of NEA0069.1 is similar to specimens identified as *Se. perturbans* from Thailand (mean interspecific p-distance 2.2%) (S2 Table), we took into account the complex history of *Se. perturbans* [88–90], and Lewis’s consideration of *Se. whartoni* as a junior synonym of *Se. perturbans* [86]. We therefore considered the specimen from Singapore as *Se. whartoni* because the identity of *Se. perturbans* remains doubtful. A photo of the specimen NEA0069.1 can be found in Fig 11.

## 5. Conclusion

To the best of our knowledge, this is the first report on the presence of phlebotomine sand flies in Singapore. This study reports four new species of sandflies along with first geographical record of one species of *Phlebotomus* and three species of *Sergentomyia* sand flies. The newly identified species are *Ph. seowpohi* n. sp., *Se. gubleri* n. sp., *Se. leechingae* n. sp., and *Se. retrocalcarae* n. sp., while the known species include *Ph. stantoni*, *Se. barraudi* group, *Se. iyengari* group, *Se. whartoni.* In this study, sandflies were found across secondary forests, public parks, peri-urban vegetated areas, open fields, and coastal areas (Fig 2 and S1 Fig). These locations represent only the current sampling sites and do not necessarily represent all areas where sand flies occur in Singapore.

While Singapore is not endemic for human leishmaniasis, its vulnerability to the disease is accentuated by its role as a major travel hub, substantial reliance on migrant workers from endemic regions, and increased travel to leishmania-endemic areas. Documented cases of imported leishmaniasis among migrant workers and returning travellers highlight the risk of introducing Leishmania and other pathogens into the country [91–93]. In 2020, antibodies of *Leishmania infantum* – which cause zoonotic leishmaniasis, was detected in a local free-roaming dog [94]. Dogs are known to be the main reservoir hosts of the disease [95–98]. This finding, coupled with risk of human leishmaniasis being introduced into Singapore and changes driven by urbanisation, land use change, and climate change [42,99–104], underscore the importance of understanding local sand fly biology, ecology diversity, distribution, and disease transmission potential. Strengthening biosurveillance efforts, by expanding surveillance sites, and conducting continuous sand flies monitoring, will help fill existing knowledge gaps and provide deeper insights into local sand fly fauna.

## Author Contributions

Huicong Ding

Majhalia Torno

Khamsing Vongphayloth

Germaine Ng

Denise Tan

Wendy Sng

Kelvin Ho

Fano José Randrianambinintsoa

Jérôme Depaquit

Cheong Huat Tan

### Conceptualization

Huicong Ding, Majhalia Torno, Jérôme Depaquit, Cheong Huat Tan

### Methodology

Huicong Ding, Majhalia Torno, Khamsing Vongphayloth, Germaine Ng, Denise Tan, Wendy Sng, Kelvin Ho, Fano José Randrianambinintsoa, Jérôme Depaquit, Cheong Huat Tan

### Resources

Huicong Ding, Majhalia Torno, Khamsing Vongphayloth, Germaine Ng, Denise Tan, Wendy Sng, Kelvin Ho, Fano José Randrianambinintsoa, Jérôme Depaquit, Cheong Huat Tan

### Supervision

Huicong Ding, Majhalia Torno, Jérôme Depaquit, Cheong Huat Tan

### Validation

Huicong Ding, Majhalia Torno, Khamsing Vongphayloth, Germaine Ng, Denise Tan, Wendy Sng, Kelvin Ho, Fano José Randrianambinintsoa, Jérôme Depaquit, Cheong Huat Tan

### Visualization

Huicong Ding, Majhalia Torno, Jérôme Depaquit, Cheong Huat Tan

### Writing – original draft

Huicong Ding, Majhalia Torno, Jérôme Depaquit, Cheong Huat Tan

### Writing – review & editing

Huicong Ding, Majhalia Torno, Khamsing Vongphayloth, Germaine Ng, Denise Tan, Wendy Sng, Kelvin Ho, Fano José Randrianambinintsoa, Jérôme Depaquit, Cheong Huat Tan

## Acknowledgements

The authors would like to express their sincere gratitude to the Malaria Surveillance Section, National Environment Agency for carrying out the entomological surveillance. We thank, Mr Chew Ming Fai, Director General for Public Health, for his support and approval to publish the study. We thank the Ministry of Finance, Singapore, for the Reinvestment Fund made available for this study.

## Supporting information Captions

**S1 Fig. Twenty-nine sampling sites across Singapore where entomological surveillance was performed in this study.** Each colored dot represents a site and its corresponding habitat type.

**S2 Fig. Maximum-likelihood *Phlebotomus* (A) and *Sergentomyia* (B) phylogenetic tree inferred from aligned consensus *cytochrome c oxidase subunit I* (*COI*) sequences using the Hasegawa-Kishino-Yano 85 model**. Sequences highlighted in red are generated from specimens collected in Singapore. Sequence in yellow is from Luangphabang, Laos, as part of IP Laos collection program. It will be further detailed in the discussion section. The trees have been rooted on reference *Sergentomyia barraudi* and *Phlebotomus stantoni* sequences which are selected as outgroups, respectively. Nodes with bootstrap value less than 70% are not shown.

**S1 Table. Mean interspecific genetic distances for *COI* sequence pairs between four *Phlebotomus* (*Euphlebotomus*) species and *Ph.* (*Anaphlebotomus*) *stantoni*. Numbers (1-7) in the top row correspond to the species listed. Calculations were based on the p-distance model. Diagonal bold values indicate intraspecific mean distances. Names with an asterisk refer to specimens collected in Singapore (SG).** NA denotes cases in which it was not possible to estimate genetic distances due to single DNA barcode.

**S2 Table. Mean interspecific genetic distances for *COI* sequence pairs between *Sergentomyia* species. Numbers (1-17) in the top row correspond to the species listed. Calculations were based on the p-distance model. Diagonal bold values indicate intraspecific mean distances. Names with an asterisk refer to specimens collected in Singapore (SG).** NA denotes cases in which it was not possible to estimate genetic distances due to single DNA barcode.

**S3 Table. Morphometric measurements (in µm) of *Phlebotomus seowpohi* n. sp. specimens.** Values are presented as mean (minimum-maximum).

## Author Contributions

Conceptualization: JD, CHT. Data curation: HD, MT, KV, GN, JD, CHT. Funding acquisition: CHT. Investigation: HD, MT, KV, GN, WS, KH, FJR, JD, CHT. Methodology: HD, MT, GN, JD, CHT. Project administration: MT, WS, KH, CHT. Resources: JD, CHT. Supervision: MT, CHT. Validation: HD, MT, JD. Visualization: HD, GN, FJR, JD. Writing – original draft preparation: HD, KV, DT, JD, CHT. Writing – review and editing: HD, DT, JD, CHT.

## Notes

### Competing Interest Statement

The authors have declared no competing interest.

